# Steering Sequence Generation in Protein Language Models through Iterative Lookback Monte Carlo Sampling

**DOI:** 10.64898/2026.05.01.722156

**Authors:** Francesco Calvanese, Gianluca Lombardi, Martin Weigt, Jorge Fernandez-de-Cossio-Diaz

## Abstract

Protein language models (pLMs) leverage large-scale evolutionary data to generate novel sequences, but steering generation toward desired physicochemical properties without sacrificing diversity remains a major challenge. Existing approaches often induce severe diversity loss or require computationally expensive retraining. We introduce **Iterative Lookback Monte Carlo** (ILMC), a training-free inference-time sampling strategy that interleaves autoregressive elongation with Metropolis–Hastings refinement to approximate sampling from a maximum-entropy target distribution balancing generative quality and steering objectives. We show theoretically that this target distribution is entropy-maximizing under fixed generative quality and steering constraints, and empirically that ILMC produces more diverse samples than standard autoregressive baselines at matched generative quality. Using simple steering potentials, ILMC improves desired molecular properties, including generating proteins with up to 12°C higher predicted melting temperature than compute-matched alternative strategies. ILMC naturally applies to classifier-guided steering, where it outperforms purely autoregressive guidance in diversity while maintaining comparable enrichment of target properties. We validate ILMC on family-specific pLMs and on the multi-family model ProGen3.

## 1. Introduction

Protein Language Models (pLMs) trained on millions of natural proteins have demonstrated a remarkable ability to capture complex evolutionary rules governing protein structure and function (Madani et al., 2023; Lin et al., 2023). These models learn a probability landscape where regions of high probability correspond to biologically active proteins. Sampling from the model can result in novel sequences whose function has been verified in experiments. Higher-probability samples are more likely to be functional in experiments (Russ et al., 2020; Fields et al., 2025), and practitioners often “steer” generation to artificially enrich for such high-probability variants (Lambert et al., 2025; Fernandez-de Cossio-Diaz et al., 2025). In textual large language models (LLMs), high-probability sampling (also called *distribution sharpening*) can elicit reasoning-like capabilities (Karan & Du, 2025), but existing methods often result in critical diversity loss (He et al., 2025; Song et al., 2025; Calvanese et al., 2025) or require computationally expensive retraining (Kim et al., 2024; Stocco et al., 2025). Furthermore, beyond sampling viable sequences, a critical challenge in protein engineering is to design specific attributes important in real-world applications, such as thermostability, or finding variants within the mutational vicinity of a wild-type sequence to preserve existing compatibilities, rather than generating totally new sequences.

To bridge the gap between unconditional generation and functional design, standard approaches rely on retraining or fine-tuning (Nijkamp et al., 2023). While effective, these methods are computationally demanding and prone to “catastrophic forgetting,” where the model over-optimizes the target property but neglects broader functional priors learned during pre-training. More recently, Reinforcement Learning (RL) has been employed to align models with desired rewards. Yet, as noted in recent literature, RL-based distribution sharpening often leads to a collapse in generation diversity (Song et al., 2025).

### Our contribution

We introduce Iterative Lookback Monte Carlo Sampling (ILMC), a training-free inference-time method for controllable generation in protein language models. ILMC targets a maximum-entropy distribution that balances the generative quality of the base model with a user-defined steering potential. Algorithmically, it interleaves standard autoregressive elongation with Metropolis– Hastings (MH) refinement steps that revisit and resample recent segments of the sequence. This correction allows ILMC to steer generation toward desired regions of sequence space without gradient updates or model retraining.

We show theoretically that the distribution targeted by ILMC is the unique maximum-entropy distribution under fixed generative quality and steering constraints. Empirically, ILMC yields more diverse samples than standard autoregressive baselines at matched generative quality. We demonstrate steering toward enhanced thermostability and toward the mutational neighborhoods of a wild-type sequence. ILMC extends naturally to classifier-guided steering for non-decomposable objectives, and we evaluate this setting on a homodimerization structural-class task. Across Chorismate Mutase, Phage Lysozyme, and Response Regulator families, and in both family-specific pLMs and ProGen3, ILMC consistently improves the diversity–quality trade-off for steered generation.

## 2. Materials and Methods

### 2.1. Generation steering and Maximum Entropy

Consider a generative model defining a probability distribution *P* (**a**) over protein sequences **a** = (*a*_1_, *a*_2_,. .., *a*_*L*_), where *a*_*i*_ ϵ *𝒜*, the set of 20 amino acids. In statistical physics terms, this model characterizes an energy landscape *E*(**a**) = log *P* (**a**). Sequences with lower energy are more probable under the model and are more likely to be functional in experiments (Russ et al., 2020; Lambert et al., 2025; Fernandez-de Cossio-Diaz et al., 2025).

Besides sampling functional variants, one may be interested in potentiating certain attributes of designed sequences, such as thermostability. Formally, we introduce a steering potential *S*(**a**), with the aim of generating sequences that minimize *S*. Importantly, *S*(**a**) is not a generative score; sequences with low *S* but high *E* may not be valid, biologically active protein sequences. Conversely, functional sequences with low *E* may have high *S*. It is therefore important to consider both scores simultaneously.

We seek a sampling strategy *Q*(**a**) satisfying **two constraints**: **i)** An expected steering score, 𝔼_*Q*_[*S*(**a**)] = *C*_S_; and **ii)** An expected generative quality (or cross-entropy) 𝔼_*Q*_[*E*(**a**)] = *C*_E_. Among all such sampling strategies, we prefer the one that remains as diverse as possible, which leads naturally to a maximum-entropy formulation:

#### Proposition 2.1

*Among all distributions Q*(**a**) *satisfying these two constraints, the one with the highest entropy* 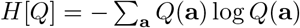 *has the form:*

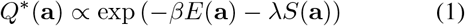

*where β and λ are Lagrange multipliers chosen to satisfy the target expectations C*_E_ *and C*_S_.

While Energy-Based Models (EBMs) are naturally suited to facilitate sampling from such distributions via Monte Carlo (MC) techniques (like Gibbs sampling or MH), applying similar methods to autoregressive models poses significant challenges (Karan & Du, 2025). An important distinction is that the inverse temperature parameter *β* can be misinterpreted in the context of the autoregressive LLMs literature, where “low-temperature sampling” often refers to the local sharpening of the next-token distribution. In that case, logits over the vocabulary of the next token are multiplied by *β*. We refer to this variant as *greedy* autoregressive low-temperature sampling. See Appendix C for further discussion on why these two approaches are fundamentally different and the computational obstacles to correctly sampling from (1). In fact, any other sampling strategy (top-*p*, beam search, etc.) (Radford et al., 2019; Holtzman et al., 2019) necessarily produces less diverse samples than *Q\**.

Given the difficulty of sampling directly from (1), it is useful to characterize how other sampling strategies *Q* relate to this optimal distribution.

#### Proposition 2.2

*For any other distributions Q*_1_(**a**), *Q*_2_(**a**) *satisfying the same constraints:*

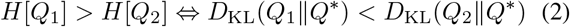

*where D*_KL_ *is the Kullback-Leibler (KL) divergence*.

See Appendices B.2 and B.1 for the proofs of 2.1 and 2.2.

In the next section we introduce ILMC, an algorithm to approximate sampling from (1). Beyond enabling attribute-targeted generation (a challenge in standard autoregressive pLMs), 2.2 ensures that approximating *Q\** (in the sense of minimizing *D*_KL_(*Q ║ Q\**)) automatically preserves sample diversity (in the sense of maximizing *H*[*Q*]).

### 2.2. Iterative Lookback Monte Carlo (ILMC)

Sampling from the base model *P* (**a**), corresponding to setting *β* = 1 and *λ* = 0 in (1), is computationally cheap in standard pLMs (Madani et al., 2023; Bhatnagar et al., 2025), which can be sampled efficiently via autoregressive decoding. ILMC leverages this base sampler to draw sequences from the tilted target *Q\** at *β >* 1 or *λ >* 0. We extend a recently proposed method (Karan & Du, 2025) to approximate sampling from distributions of the form ∝ *P* (**a**)^*β*^, a particular case of (1) with *λ* = 0, *i*.*e*., without steering. Here we extend this framework to incorporate a steering potential *S*(**a**).

To describe our sampling algorithm, we consider an intermediate iteration where a sequence prefix **a**_*≤t*_ = (*a*_1_, *…, a*_*t*_) has already been generated. We introduce a partial generative energy and steering potential defined on the prefix:

- **Partial Generative Energy** *E*_*t*_(**a**≤_*t*_), defined as the cumulative negative log-likelihood of the prefix under the base generative model:

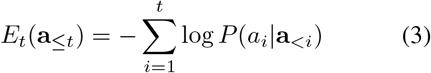
- **Partial Steering Potential** *S*_*t*_(**a**≤_*t*_), which depends on the attribute one aims to optimize. It must satisfy two formal requirements: i) must not depend on future tokens, **a**_>*t*_; and ii) must be compatible with the steering score at end-of-sequence, *i*.*e*., *S*_*L*_(**a**_*L*_)= *S*(**a**).

With these quantities defined, the procedure to sample the rest of the sequence consists of alternating Elongation and Refinement steps. The algorithm depends on three input parameter settings: block-size *B*, lookback window *K*, and number of MC steps *N*_MC_.

**Step 1: Autoregressive Elongation**. First, we extend the current prefix by sampling a block of *B* new tokens. For each position *i* ϵ*{t* + 1,. .., *t* + *B}*, draw

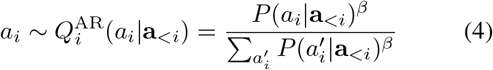

as in greedy “low-temperature” autoregressive sampling. This process is repeated *B* times to extend the sequence, after which the position index is updated to *t* ← *t* + *B*.

**Step 2: Lookback Refinement and Steering**. To correct the distribution towards the target potential defined by β*E*_*t*_(**a**_≤*t*_) + *λS*_*t*_(**a**_≤*t*_), we perform *N*_MC_ steps of a Metropolis-Hastings refinement on the current sequence segment. In each refinement step:

1. **Selection**: A pivot position *j* is chosen at random from the lookback window *{t* − *K* + 1,.. ., *t}*.
2. **Resampling**: A candidate variant 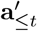 is generated. The prefix 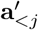 remains identical to the current sequence **a**_*<j*_. The suffix from *j* to *t* is resampled autoregressively using the proposal distribution *q* defined in Eq. (4), until the length *t* is reached.
3. **Acceptance**: The candidate 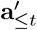 is accepted with a MH probability:

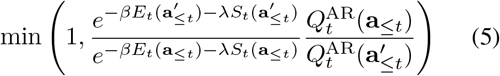

The proposal distribution 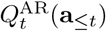 is the product of the autoregressive steps used to generate the sequence up to *t*:

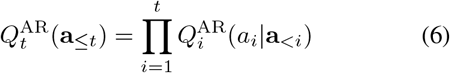

Note that in the ratio 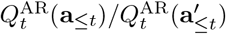, terms *i < j* cancel because of the shared prefix up to *j*.

This procedure repeats until the end-of-sequence token (⟨EOS⟩) is reached. While the complete sequence energy *E*(**a**) and steering potential *S*(**a**) are only fully realized at the final iteration, the intermediate *S*_*t*_(**a**_≤*t*_) and *E*_*t*_(**a**_≤*t*_) steer the chain towards high-probability regions early in the generation process.

When *K* = 1, ILMC mathematically reduces to greedy autoregressive low-temperature sampling. In the idealized limit *N*_MC_ → ∞ and full lookback *K* = *L*, the procedure converges to the target distribution *Q*^*^(**a**).

In practice, a restricted lookback window *K < L* and moderate *N*_MC_ are sufficient to reach high-quality samples in realistic pLMs, as we show in Sec. 3. Moreover, a smaller *K* results in higher acceptance rate of the proposals made by *Q*^AR^ (4), which lacks steering. If *K* were large, the proposals sample long segments without guidance from *S*_*t*_(**a**_≤*t*_). The probability that such non-steered segments have good steering scores can be low. Consequently, we observe a decrease in acceptance probability as a function of *K*. See Appendix L for more discussion of this trade-off and Appendix P for guidelines on choosing these hyperparameters.

### 2.3. Effective steering potentials

ILMC depends critically on the choice of the steering potential *S*_*t*_(**a**_≤*t*_). While any sequence-level score can in principle define a target distribution *Q*^*^, the autoregressive nature of our sampler requires that the corresponding partial potential provides a meaningful signal early in the generation process.

Importantly, a full-sequence steering objective *S*(**a**) does not uniquely determine its prefix decomposition *S*_*t*_(**a**_≤*t*_). In addition to choosing the objective itself, one must choose how to distribute it along the generation process. Effective decompositions make the dependence on the current token as explicit as possible while minimizing residual dependence on earlier tokens. As discussed in Appendix F, strong residual dependence on the past leads to a larger mismatch between the practical sampler and the ideal target *Q*^*^, although larger lookback windows *K* can partially compensate for this effect.

In this work, we consider two kinds of steering signal constructs. First, we use objectives with natural prefix decompositions: an additive composition-based score for thermostability and a prefix-compatible distance function for wild-type proximity. Second, for the more challenging non-decomposable objective of homodimerization structural class, we show that ILMC can be naturally extended to classifier-guided steering. In both settings, the steering signal biases sampling toward the desired region of sequence space, while the base pLM preserves generative quality and suppresses unrealistic or non-functional sequences.

Additional examples, mathematical details, and a broader discussion of effective and ineffective steering potentials are provided in Appendices E and F.

### 2.4. Computational complexity and scalability

ILMC sampling introduces a refinement phase consisting of *N*_MC_ MH steps after each elongation block of size *B*. As derived in Appendix G, the total computational cost scales approximately as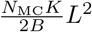, assuming that the main computational bottleneck is the evaluation of the generative score *E*_*t*_(**a**_≤*t*_) rather than the steering score *S*_*t*_(**a**_≤*t*_), whose cost depends on its specific form and is not the limiting factor in the settings studied here. This relation highlights the dependence on the lookback window *K*. When the lookback window is non-extensive (fixed *K*, independent of *L*), ILMC remains in the same asymptotic complexity class as standard autoregressive sampling with key-value caching, namely *O*(*L*^2^), albeit with a larger constant factor. By contrast, when the lookback window is extensive (*K* ∝ *L*), the complexity scales as *O*(*L*^3^).

## 3. Results and Discussion

To validate ILMC, we consider three protein families: **Chorismate Mutase (CM), Phage Lysozyme (PL)**, and **Response Regulator (RR)**. CM and PL are well-established benchmarks where generative models have successfully produced biologically functional artificial variants (Russ et al., 2020; Madani et al., 2023). RR is used to evaluate classifier-guided steering on a harder structural objective, and connects our setting to recent pLM steering literature (Caredda et al., 2026). We evaluate ILMC across two architectures of different scales: 1) **Small Decoder-Only (sDO)**, a compact autoregressive Transformer with approximately 0.8M parameters, trained from scratch on the natural sequences of each family; 2) **ProGen3**, a 112M-parameter version of ProGen3 (Madani et al., 2023; Bhatnagar et al., 2025), the large-scale pLM pre-trained on UniProt (The UniProt Consortium, 2025) and subsequently fine-tuned on the specific family datasets. See Appendix H for details on the architectures, training data, optimization protocols, and model hyperparameters.

### 3.1. Generating Diverse High-Probability Samples

We first investigate the case where no steering potential is applied, i.e., *S*(**a**)= 0. From Proposition 2.1, sampling from the exponentiated distribution 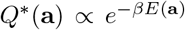is the unique strategy that maximizes entropy for a fixed expected generative quality *C*_E_. In an Energy-Entropy plane, this defines a Pareto efficiency frontier; for any given mean energy 𝔼_*Q*_[*E*], no sampling strategy can achieve higher diversity than *Q*^*^. Conversely, *Q*^*^ reaches the lowest possible average energy for a fixed entropy budget.

Empirically, the lookback window *K* is a critical determinant of the quality-diversity trade-off. To achieve a specific target mean energy, standard autoregressive sampling (*K* = 1) requires a significantly higher inverse temperature *β* than ILMC with *K >* 1, and the required *β* decreases as *K* increases (see Table 3). This high local sharpening in the autoregressive case collapses the distribution’s support, leading to a marked reduction in diversity. Using the number of unique draws over a sample of 1024 sequences as a proxy for entropy, we find that increasing *K* lets ILMC reach high-probability sequences while maintaining a significantly higher count of unique variants, as illustrated in Figure 2 (sDO model, PL family). While a higher *K* intuitively provides a better approximation of the target *Q*\*, and thus higher entropy per Proposition 2.2, this is not always mathematically guaranteed, as it is possible to find pathological cases that defy this intuition; see Appendix D. Furthermore, because we set a fixed number of MH steps (*N*_MC_ = 10) and do not reach strict equilibrium, even when *K* = *L* the sampling only approximates *Q*\*.

**Figure 1.**
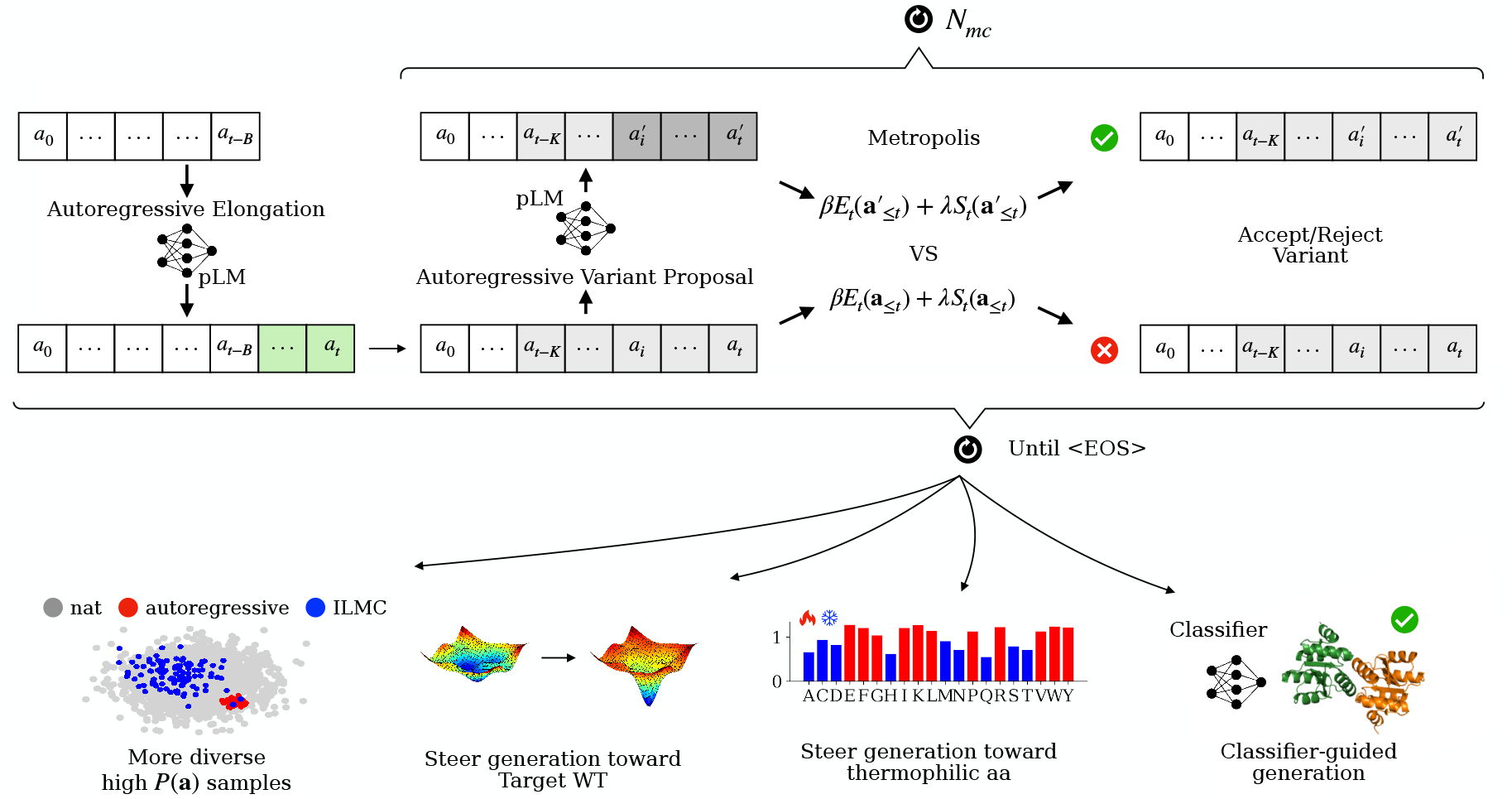
**Iterative Lookback Monte Carlo (ILMC)** sampling framework. The algorithm approximates sampling from the target *Q*^*\**^ (**a**), (1), by interleaving two phases: Autoregressive Elongation and MH Refinement. In the elongation phase, the pLM extends the sequence prefix by a block of tokens. In the refinement phase, a variant is proposed by resampling a segment of the generated history using the pLM; this variant is accepted or rejected based on a global potential that combines the generative energy *E* of the model and a user-defined steering potential *S*. This procedure promotes higher sampling diversity and enables the steering of the generation toward regions of the sequence space that satisfy specific properties, such as thermostability or vicinity to a target wild-type, without requiring gradient updates or model retraining.

**Figure 2.**
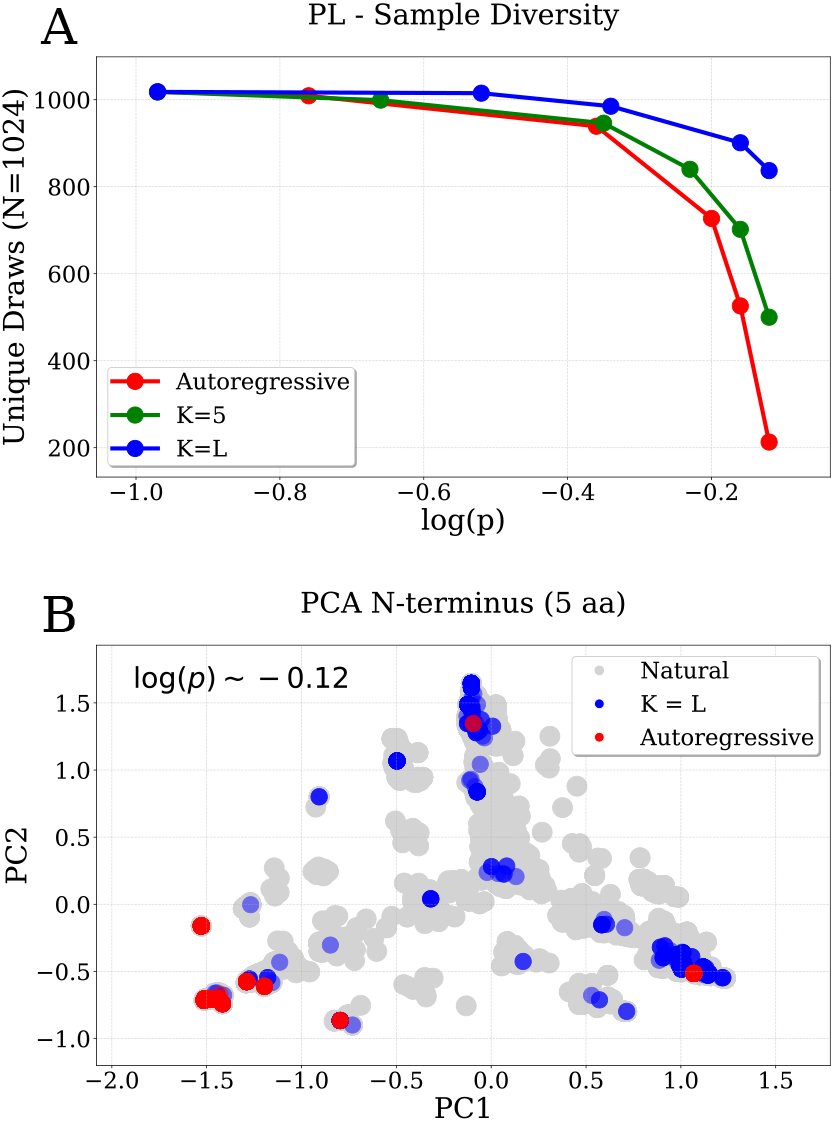
**Diversity vs. generative quality** (sDO on PL, 1024 sequences per sample). (A) Unique sequences (*N* = 1024) vs. log *P*. Standard autoregressive sampling (red) suffers from rapid diversity collapse, while ILMC (*K* = 5 green, *K* = *L* blue; *B* = 2, *N*_MC_ = 10) draws more unique variants. (B) PCA of N-termini (first 5 residues) at high probability (log *P* ≈ −0.12). Standard sampling (red) collapses into narrow modes, while global lookback (*K* = *L*, blue) covers the natural sequence space (grey).

Various library diversity metrics across sDO and ProGen3 models and the two families (PL and CM), using sampling parameters *B* = 2, *N*_MC_ = 10 and varying *K* and *β*, are reported in Table 3. Specifically, we track the total number of unique draws (out of 1024), the number of unique draws within an edit distance radius of 5 (meaning that once a sequence is selected, all subsequent draws with a distance ≤ 5 are considered duplicates), the mean pairwise distance within the library (Dist Gen-Gen), and the mean edit distance to the closest neighbor (Min Dist Gen-Gen). These metrics generally improve as *K* increases, in agreement with Proposition 2.2. The results consistently show increased diversity for both sDO and ProGen3 across both families, indicating that the method scales effectively. In contrast, autoregressive samples exhibit a pronounced mode collapse at high log *P*, evident in the histogram of pairwise distances (Dist Gen-Gen), shown in Appendix Fig. 6.

This high-log *P* regime, where ILMC yields its largest diversity gains, is particularly relevant in practical protein design. In downstream experimental pipelines, practitioners often prioritize the highest-scoring sequences through highly selective strategies such as greedy decoding (*β ⟶ ∞* limit) (Faizi et al., 2025) or post-hoc filtering (Lambert et al., 2025), because high-likelihood proteins are more likely to remain functional in wet-lab validation (Russ et al., 2020; Fields et al., 2025).

Visual inspection of the sampled libraries reveals that increasing the lookback window *K* leads to a significantly wider distribution of sequence variety at the N-terminus in the high log *P* samples. We show this in Figure 2 for the sDO model applied to the PL family, which presents a Principal Component Analysis (PCA) projection of the first five amino acids, comparing standard autoregressive sampling (*K* = 1) with the global lookback case (*K* = *L*). Additional examples are provided in Appendix Fig. 7. This phenomenon can be intuitively attributed to the inverse temperature *β*. In the *K* = 1 regime, achieving a given low-energy target requires very high *β* (see Table 3), which effectively collapses the initial generation to few sequences as the local logit profile becomes very sharp. Diversity is only introduced when a lower-probability token, or “stochastic error,” is sampled, but as *β* increases, the waiting time for this “error” event grows, causing a collapse in the diversity of the N-terminus. In contrast, ILMC allows the model to explore a broader set of initial states, that are later refined into high-quality sequences. This also suggests that the high-probability samples at higher *K* settings better cover the sequence space around the natural sequences. To quantify this, Table 3 reports the mean distance of natural sequences to a closest sequence of the generated library (Min Dist Gen-Nat). Lower values indicate that the generated samples better explore the region of sequence space occupied by natural proteins. This metric tends to decrease with *K* at a given generative quality (mean value of log *P*), confirming that the samples explore a broader region of the sequence space. A comparison with top-*p* sample diversity is provided in the Table 4.

### 3.2. Steering generation toward enhanced thermostability

Comparative proteomic analyses have shown that amino acid composition is a primary determinant of thermal stability. Specifically, thermophilic organisms exhibit a distinct enrichment of a subset of residues, often identified by the mnemonic IVYWREL, which facilitate the formation of robust salt bridges, improved hydrophobic packing, and increased hydrogen bonding (Fukuchi & Nishikawa, 2001; Zeldovich et al., 2007). In this experiment, we define a steering potential *S*^Tm^(**a**) to bias generation toward these stabilizing amino-acids. Following established findings that surface residues are less constrained by evolutionary conservation and more responsive to environmental adaptation, we use propensity scores derived specifically from surface amino acid compositions (Fukuchi & Nishikawa, 2001).

For an amino acid *a ∈ 𝒜*, let *w*(*a*) denote the ratio of its frequency in thermophilic surfaces to its frequency in mesophilic surfaces. The table of propensities *w*(*a*) is provided in Appendix J. We define the global steering potential as the negative sum of these ratios across the sequence:

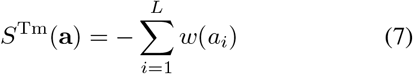

Optimizing this score in isolation would not yield meaningful protein designs: one could trivially increase it by enriching sequences in favorable residues without preserving global plausibility or function. ILMC avoids this failure mode by balancing the steering score against the generative energy *E*(**a**). Steering therefore operates only within regions of sequence space that remain plausible under the pLM. In this sense, ILMC can turn simple property-correlated proxy scores into practically useful design objectives.

The partial steering potential for an incomplete prefix **a**_≤*t*_ is naturally defined as the cumulative sum over the existing residues:

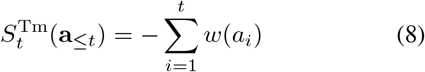

This additive formulation ensures that the criteria for effective steering discussed in Section 2.2 are satisfied, providing a steering signal from the initiation of the sequence.

The global target distribution is then (*c*.*f*. Eq. (1)):

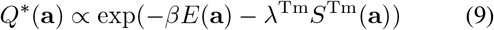

where *λ*^Tm^ is the Lagrange multiplier associated with the steering potential *S*^Tm^(**a**). Interestingly, we observe that increasing *λ*^Tm^ at constant *β*, decreases the average *S*^Tm^ but *increases* the average generative *E*. To maintain a target generative quality, we must also increase *β*. This indicates an inherent trade-off between the generative energy and thermostability for the designed sequences. These results are reported in Table 1. Further analysis regarding the Pareto frontier between generative energy and the steering score are provided in Appendix M.

**Table 1.** Performance of Thermostability Steering. We compare standard autoregressive sampling (Base: λ^Tm^ = 0), a compute-matched rejection-sampling baseline (Rejection), and ILMC (Steered: λ^Tm^ = 20,*B* = 2,*K* = 5, *N*_MC_ = 10) across two architectures (sDO, ProGen3) and protein families (CM, PL). Statistics are computed over 1024 sequences per sample. Metrics include: log *P* (mean per-token log *P*); pLDDT (predicted local structural confidence via ESMFold); *T*_*m*_ (predicted melting temperature via TemStaPro); Dist Gen-Gen (mean pairwise edit distance within the library); Min Dist Gen-Gen (mean edit distance to the nearest generated neighbor); and Min Dist Gen-Nat (mean edit distance to the nearest natural sequence).

| FAMILY | MODEL | STRATEGY | $\beta$ | $\log P$ | pLDDT | Tm<br>(°C) | LENGTH | DIST<br>(GEN-GEN) | MIN DIST<br>(GEN-GEN) | MIN DIST<br>(GEN-NAT) |
| --- | --- | --- | --- | --- | --- | --- | --- | --- | --- | --- |
| CM | sDO | BASE | 1.00 | -0.80 ± 0.46 | 78 ± 4 | 41.1 ± 6.5 | 89.8 ± 1.6 | 64.5 ± 9.8 | 29.3 ± 16.9 | 25.7 ± 16.8 |
|  |  | REJECTION | 1.50 | -0.82 ± 0.28 | 78 ± 4 | 56.1 ± 11.0 | 89.4 ± 1.7 | 59.8 ± 8.5 | 30.6 ± 12.1 | 33.0 ± 13.4 |
|  |  | STEERED | 1.80 | -0.78 ± 0.31 | 78 ± 4 | **61.5 ± 10.8** | 90.0 ± 1.8 | 53.1 ± 14.0 | 20.8 ± 12.4 | 35.0 ± 17.1 |
|  | ProGen3 | BASE | 1.10 | -0.62 ± 0.45 | 79 ± 3 | 40.8 ± 6.4 | 88.6 ± 5.8 | 63.8 ± 9.9 | 25.9 ± 15.0 | 22.8 ± 15.8 |
|  |  | REJECTION | 1.80 | -0.63 ± 0.27 | 79 ± 2 | 55.6 ± 11.6 | 89.6 ± 2.2 | 60.0 ± 10.0 | 27.4 ± 14.1 | 30.2 ± 14.7 |
|  |  | STEERED | 2.00 | -0.60 ± 0.25 | 79 ± 2 | **60.8 ± 11.0** | 89.3 ± 1.8 | 56.4 ± 10.2 | 21.5 ± 11.5 | 29.7 ± 14.9 |
| PL | sDO | BASE | 1.00 | -0.97 ± 0.63 | 60 ± 11 | 39.2 ± 5.6 | 128.1 ± 5.9 | 91.2 ± 13.0 | 48.7 ± 28.6 | 45.4 ± 28.7 |
|  |  | REJECTION | 1.10 | -1.06 ± 0.63 | 57 ± 10 | 45.5 ± 11.3 | 127.6 ± 6.7 | 92.5 ± 13.9 | 52.5 ± 28.4 | 51.5 ± 28.1 |
|  |  | STEERED | 1.30 | -0.96 ± 0.54 | 59 ± 11 | **57.3 ± 13.7** | 132.4 ± 5.3 | 89.4 ± 16.9 | 45.0 ± 27.1 | 51.7 ± 29.5 |
|  | ProGen3 | BASE | 1.20 | -0.53 ± 0.40 | 67 ± 7 | 39.7 ± 6.6 | 128.2 ± 6.9 | 85.6 ± 15.0 | 36.1 ± 26.0 | 34.0 ± 25.7 |
|  |  | REJECTION | 1.60 | -0.53 ± 0.30 | 64 ± 8 | 48.4 ± 12.8 | 128.1 ± 7.3 | 81.1 ± 17.4 | 33.9 ± 22.7 | 37.2 ± 23.3 |
|  |  | STEERED | 1.50 | -0.52 ± 0.32 | 67 ± 7 | **54.4 ± 13.9** | 130.6 ± 13.4 | 85.5 ± 24.4 | 30.8 ± 24.0 | 38.7 ± 30.2 |

To assess whether this sequence-level steering translates into thermophilic adaptation, we predicted melting temperatures (*T*_*m*_) of generated sequences using TemStaPro (Pudžiuvelytė et al., 2024) (implementation details in Appendix I). At matched generative quality, ILMC increases the predicted *T*_*m*_ by an average of 18.3^*°*^C relative to the standard autoregressive baseline and 7.1^*°*^C against a runtime-matched rejection-sampling obtained by generating standard autoregressive samples with the same total compute budget and filtering them by *S*^Tm^(**a**). To verify that steering does not degrade structural plausibility, we also computed pLDDT scores with ESMFold (Lin et al., 2023), finding that the optimization of *S*(a) does not measurably reduce predicted structural confidence.

Results for the sDO model on the PL family are illustrated in Figure 3, and quantitative results for both sDO and ProGen3 are reported in Table 1. Sampling parameters were set to *B* = 2, *K* = 5, and *N*_MC_ = 10, with varying *β* and *λ*^Tm^. ILMC thus steers generation toward thermostable regions of sequence space that are inaccessible to standard sampling while avoiding diversity collapse, as shown by the library-distance metrics reported in Table 1 (Dist Gen-Gen, Min Dist Gen-Gen, and Min Dist Gen-Nat).

**Figure 3.**
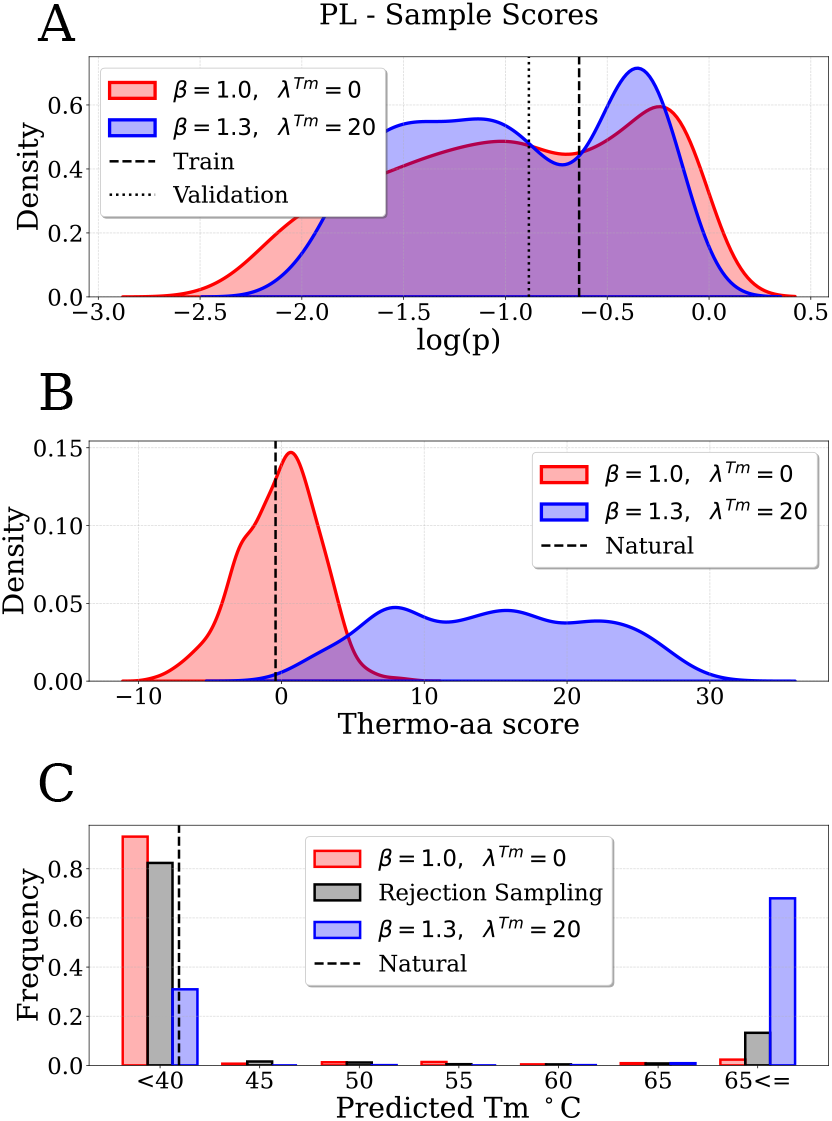
**Steering generation toward thermophilic adaptation** (sDO on PL, 1024 sequences per sample). (A) Distribution of log *P* comparing the unsteered baseline (red) with the steered ILMC sampling strategy (blue, λ^Tm^ = 20, *β*= 1.3,*B* = 2,*K* = 5, *N*_MC_ = 10). The parameters of the ILMC were explicitly tuned to match the log *P* of the baseline. (B) Distribution of the thermophilic amino acid score −*S*^Tm^(**a**). The steered samples exhibit a significant shift toward higher scores, far exceeding the range observed in the unsteered baseline and the natural sequences (dashed line). (C) Predicted melting temperatures (*T*_*m*_) estimated using TemStaPro.

### 3.3. Steering generation towards the vicinity of a target wild-type

Many applications in biological sequence engineering require mutating an existing wild-type (WT) sequence rather than generating *de novo* proteins (Calvanese et al., 2025; Lambert et al., 2025; Hie et al., 2024; Blalock et al., 2025). Identifying low-energy states in the local vicinity of natural variants is of significant biological and engineering relevance. In this section, we show that by introducing an appropriate steering potential, we are able to bias the presented sampling process toward the mutational neighborhood of a specific target.

While the edit distance is a natural metric for string divergence, it turns out to be a poor steering potential in terms of the criteria discussed in Section 2.2. For example, the raw edit distance between a low-length prefix and a full-length target is mostly determined by the length difference and is almost independent of the prefix compatibility with the beginning of the target. To address this, we define a modified edit distance. Let a_≤*t*_ be the sequence prefix generated so far, and w the target wild-type of length *L*_*w*_. We define *S*^WT^(a_*≤t*_) as the minimum number of edits required to transform the prefix a_*≤t*_ into any prefix of the target w:

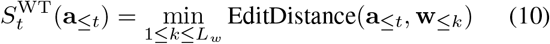

The complete target distribution for the steered sampling process is:

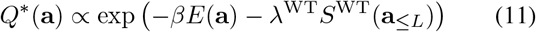

where *λ*^WT^ is the Lagrange multiplier tuning the distance towards w. This formulation ensures that 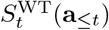provides a meaningful steering signal from the beginning of the generated sequence. Details on the implementation of this distance, along with illustrative examples and more discussion justifying its steering effectiveness, are provided in Appendix K.

Our results demonstrate that by tuning the steering strength *λ*^WT^, we can sample variants closer to the target wild-type (see Figure 4A). The generative energies of these proximal designs are distributed around the energy of the original wild-type and are significantly lower than those of random mutations at the same distance, as shown in Figure 4B (illustrated for sDO on PL). Table 5 reports the average log *P* and edit distance to the target wild-type of steered samples for both protein families and model architectures. The chosen target wild-type was the first in the provided training dataset (index 0). Sampling parameters were set to *B* = 2,*K* = 5, and *N*_MC_ = 300. Runtime-matched rejection sampling systematically fails to reach the desired mutational neighborhood: as seen in Figure 4A, the tail of the standard autoregressive distribution remains far from the target WT, so sampling close variants by post-hoc filtering would require prohibitively large sample sizes.

**Figure 4.**
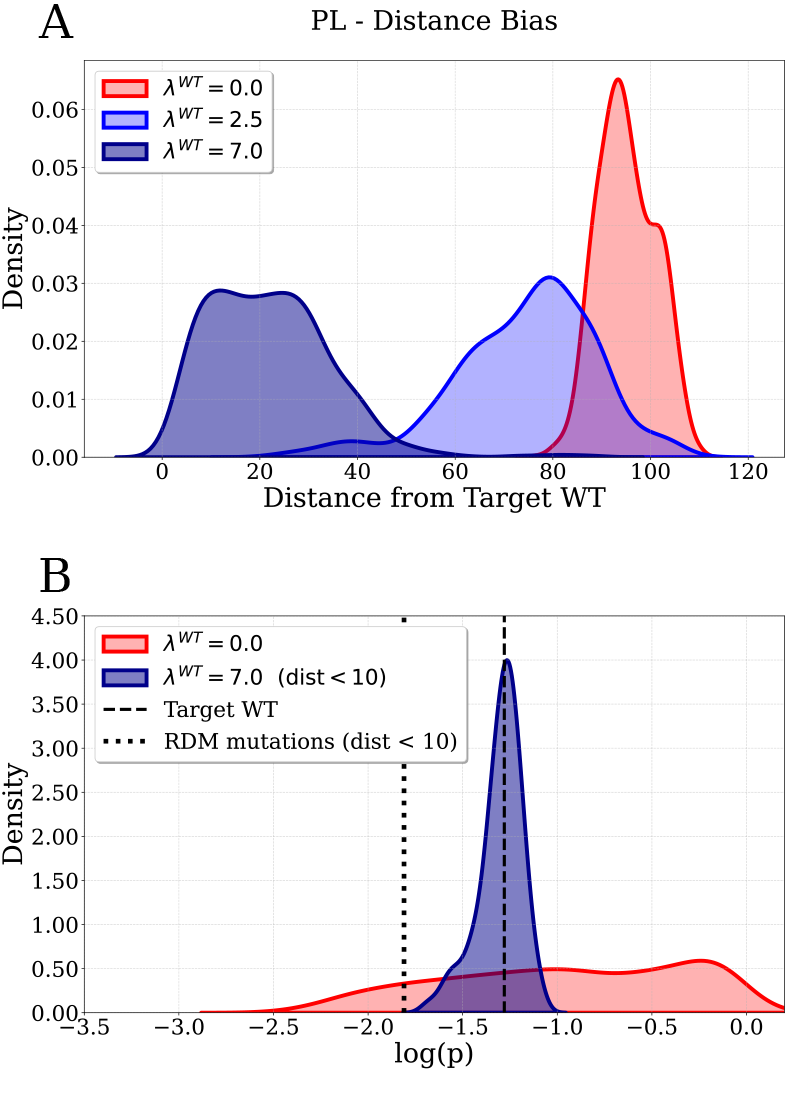
**Steering toward a target wild-type** (sDO on PL, 1024 sequences per sample). (A) Edit distance distributions for increasing steering strength λ^WT^ (*B* = 2,*K* = 5, *N*_MC_ = 300). Higher λ^WT^ concentrates sampling in the immediate local neighborhood (dark blue). (B) log *P* of variants within 10 mutations. Steered samples cluster around the WT score (dashed) and significantly outperform random mutations (dotted) compared to the unsteered baseline (red).

Another interesting application of this framework is multi-objective design, achieved by introducing two steering potentials simultaneously. For example, we could aim to enhance thermostability (*T*_*m*_) starting from a given wild-type sequence, while restricting the number of modifications on the original sequence. This task is naturally addressed with ILMC, by combining the steering potentials for thermostability and wild-type vicinity, Equations (8) and (10). ILMC will then search for thermostable variants within a small mutational radius of a wild-type target. Our results (Appendix N) confirm that ILMC is able to identify high-likelihood sequences in the close vicinity of the given wild-type with significantly improved predicted thermostability.

### 3.4. Classifier-guided steering for non-decomposable properties

The steering potentials considered so far admit natural prefix decompositions, making them particularly suitable for ILMC. However, many important protein-design objectives are not easily expressible as additive or otherwise decomposable sequence scores. Examples include global structural classes and complex functional labels that arise from long-range sequence constraints. To extend ILMC to this setting, we combine it with *classifier-guided steering*. This also enables a direct comparison with classifier-guided autoregressive decoding methods such as FUDGE (Future Discriminators for Generation) (Yang & Klein, 2021), of which ILMC can be viewed as a non-myopic extension.

FUDGE trains an any-prefix classifier, i.e., a predictor of the target label from an incomplete prefix, *p*_*ϕ*_ (*y* |a_≤*t*_), and uses it to bias next-token generation during autoregressive decoding. Intuitively, prefixes that remain compatible with multiple classes receive only weak guidance, whereas continuations that make the target class unlikely are down-weighted immediately. Concretely, if *y* denotes the target label, FUDGE samples each token from

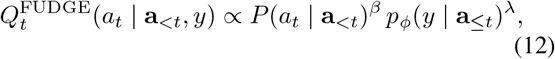

and the probability of a full sequence factorizes as

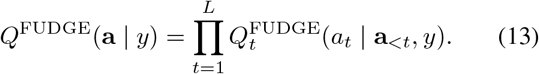

As with greedy low-temperature sampling, this construction is purely local and therefore myopic.

Within ILMC, the same prefix classifier is used differently. Rather than sampling directly from the locally guided autoregressive product in Eq. (13), we use the classifier output to define a global target distribution and approximate sampling from it with lookback Monte Carlo refinement. Specifically, we define the prefix steering potential

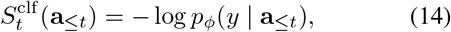

which yields the target distribution

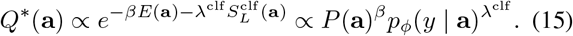

The Response Regulator (RR) receiver-domain family (Pfam PF00072 (Mistry et al., 2021)) consists of different subclasses that are associated with distinct homodimerization geometries (Gao & Stock, 2009). We train an autoregressive sDO model on the RR multiple-sequence alignment (*L* = 111). We then use a lightweight any-prefix classifier (see Appendix O) to steer generation toward a specific GerE homodimerization target (Caredda et al., 2026). Since the generator and prefix classifier are identical across the FUDGE and ILMC conditions, the comparison isolates the effect of the *sampling strategy*, rather than differences in the models.

Empirically, both FUDGE and ILMC (*B* = 2, *K* = 10, *N*_*MC*_ = 10) enrich for the target RR structural class, as assessed by AlphaFold3 (Abramson et al., 2024) based validation against the reference dimerization geometry (Table 2; Fig. 5E). At moderate sharpening (log *P* = −0.56), the two methods produce a similarly large number of unique sequences within edit-distance radius 5 (1000 for FUDGE and 1004 for ILMC), but ILMC already preserves higher intra-library diversity, with mean pairwise edit distance 63.9 ±13.3 compared with 58.35 ±13.7 for FUDGE (Table 2). Under stronger sharpening (log *P* = −0.37), the diversity advantage increases: ILMC produces 927 unique sequences within radius 5, versus 709 for FUDGE, and maintains substantially broader intra-library diversity (63.4 ±15.1 versus 48.8± 16.4; Table 2). This broader exploration is also visible in sequence space (Fig. 5C and D). Both methods strongly improve over the unsteered baseline in terms of iRMSD, but ILMC more closely spans the iRMSD range observed for natural sequences from the correct structural class (Table 2; Fig. 5E), with values consistent with the existing literature (Caredda et al., 2026).

**Table 2.** Classifier-guided steering on Response Regulators. Comparison of unsteered autoregressive sampling, FUDGE, and ILMC on the RR benchmark, reported at two matched mean pertoken log *P* operating points. Metrics are Unique (Radius 5), Dist Gen-Gen, and iRMSD (AlphaFold3) to the target structure (GerE, PDB 4ZMS). For reference, natural sequences in this structural class have an iRMSD of 11.3 *±* 3.4, vs. 15.1 *±* 4.4 for sequences from other structural classes.

| $\log P$ | METHOD | $\beta$ | UNIQUE<br>(RADIUS 5) | LENGTH | DIST<br>(GEN-GEN) | iRMSD<br>(Å) |
| --- | --- | --- | --- | --- | --- | --- |
| -0.56 | BASE | 2.0 | 1006 | 111 | $67.7 \pm 10.9$ | $17.3 \pm 3.5$ |
| | FUDGE | 1.8 | 1000 | 111 | $58.35 \pm 13.7$ | $9.8 \pm 4.3$ |
| | ILMC | 1.4 | 1004 | 111 | $63.9 \pm 13.3$ | $10.6 \pm 6.5$ |
| -0.37 | BASE | 3.5 | 694 | 111 | $63.9 \pm 15.3$ | $15.0 \pm 5.0$ |
| | FUDGE | 2.8 | 709 | 111 | $48.8 \pm 16.4$ | $9.0 \pm 3.8$ |
| | ILMC | 1.8 | 927 | 111 | $63.4 \pm 15.1$ | $10.7 \pm 3.8$ |

**Figure 5.**
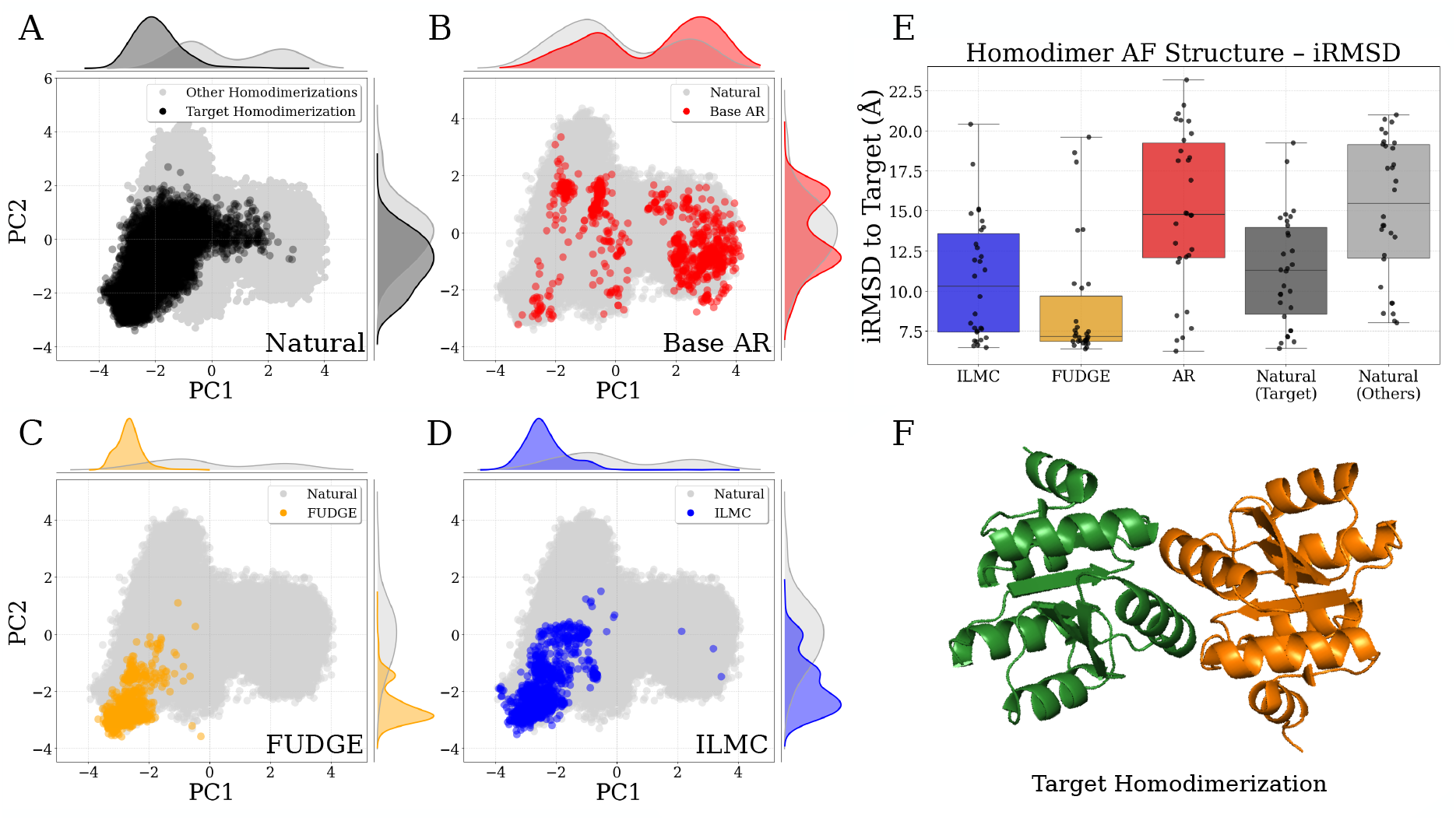
Classifier-guided steering on the Response Regulator family. All generated samples are shown at matched mean per-token log *P* = -0.37. (A) PCA projection of natural sequences (*N* = 855,395), highlighting sequences with the target GerE homodimerization class in black. (B-D) 1024 samples from unsteered autoregressive, FUDGE, and ILMC. (E) iRMSD to target GerE class (*N* = 30). (F) GerE target homodimerization structure (PDB 4ZMS).

Increasing the lookback window yields diminishing diversity gains. For example, *K* = 20 yields modest improvements (Unique within radius 5: 914; mean intra-library distance: 65.8 *±* 14.4).

These results show that ILMC is not restricted to simple decomposable steering scores. When no such proxy is available, a lightweight prefix classifier can be used as a modular steering potential, extending ILMC to non-decomposable properties while retaining a clear advantage over purely autoregressive guidance in library diversity.

## 4. Conclusion

ILMC is a *drop-in* sampler for steering pLM generation, mitigating the diversity collapse often seen with probability sharpening and reward-based decoding. By targeting a maximum-entropy distribution that balances generative likelihood with steering constraints, it produces diverse, high-quality libraries. ILMC can turn simple property-correlated proxies into effective design objectives, and can also be applied to classifier-guided steering, acting as a non-myopic correction to purely autoregressive methods such as FUDGE. From family-specific models to the state-of-the-art ProGen3, ILMC generates diverse, high-likelihood sequences and supports practical steering signals such as thermostability, wild-type proximity, and homodimerization structural class. ILMC can be broadly useful for protein engineering, with natural extensions to richer multi-objective steering and experimental validation.

## Data and Code Availability

The code associated with this paper is available in the GitHub repository: https://github.com/FrancescoCalvanese/pLM-steer-ILMC.

The data, including model checkpoints, natural protein sequences, generated protein sequences, and related analysis scripts, are available on Zenodo: https://doi.org/10.5281/zenodo.19866140.

## Funding

The authors acknowledge funding from the AISSAI Centre, funded by Google DeepMind and the CNRS Foundation. The authors also acknowledge Sorbonne Université and Institut de Physique Théorique, Université Paris-Saclay, CEA, for computational resources.

## Author Contributions

Conceptualization: F.C., G.L., M.W., J.F.-d.-C.-D. Code and analysis: F.C., G.L. Manuscript writing: F.C., J.F.-d.-C.-D. Project supervision: M.W., J.F.-d.-C.-D. Interpretation of results: F.C., G.L., M.W., J.F.-d.-C.-D. All authors reviewed and edited the manuscript.

## Conflict of Interest

The authors declare no competing interests.

## Ethical Statement

This study is computational and did not involve human participants, human data, animal experiments, or clinical samples. The generated protein sequences belong to non-harmful protein families and were used for methodological evaluation only; they were not experimentally synthesized or tested as part of this work.

## Notes

### Competing Interest Statement

The authors have declared no competing interest.

