## Appendix for "Steering Sequence Generation in Protein Language Models through Iterative Lookback Monte Carlo Sampling"

### A. Supplementary results

*Table 3.* **Diversity metrics for unsteered sampling strategies.** We compare standard autoregressive sampling (Base, $K = 1$) with ILMC ($B = 2, N_{\mathrm{MC}} = 10, K = 5, L$). All sample sets consist of 1024 sequences. Reference metrics for Natural Datasets: for CM, mean pairwise dist. $65.2 \pm 9.1$, min dist. $11 \pm 12$; for PL, mean pairwise dist. $91.3 \pm 12.2$, min dist. $17.4 \pm 20.0$.

| FAMILY | MODEL | STRATEGY | $\beta$ | $\log P$ | UNIQUE (ON 1024) | UNIQUE (RADIUS 5) | LENGTH | DIST (GEN-GEN) | MIN DIST (GEN-GEN) | MIN DIST (GEN-NAT) | MIN DIST (NAT-GEN) |
|---|---|---|---|---|---|---|---|---|---|---|---|
| **CM** | PROGEN3 | BASE | 1.30 | -0.49 ± 0.29 | 1021 | 963 | 90.0 ± 1.50 | 63.7 ± 10.8 | 22.1 ± 14.8 | 19.1 ± 14.5 | 26.9 ± 16.2 |
| | | $K = 5$ | 1.20 | -0.48 ± 0.29 | 1022 | 964 | 89.4 ± 2.0 | 63.4 ± 11.0 | 21.2 ± 14.5 | 18.2 ± 14.4 | 26.9 ± 16.2 |
| | | $K = L$ | 1.04 | -0.47 ± 0.27 | 1022 | 973 | 88.2 ± 1.7 | 60.7 ± 12.8 | 18.9 ± 13.5 | 16.2 ± 12.8 | 28.7 ± 16.3 |
| | | BASE | 2.00 | -0.31 ± 0.19 | 961 | 793 | 90.0 ± 1.5 | 63.1 ± 12.9 | 13.1 ± 13.0 | 12.2 ± 11.8 | 26.6 ± 16.2 |
| | | $K = 5$ | 1.50 | -0.30 ± 0.18 | 982 | 808 | 89.8 ± 1.7 | 63.2 ± 12.0 | 12.7 ± 12.2 | 10.8 ± 11.3 | 26.0 ± 16.7 |
| | | $K = L$ | 1.08 | -0.31 ± 0.16 | 1009 | 862 | 88.8 ± 1.4 | 58.2 ± 15.7 | 11.2 ± 9.6 | 9.0 ± 7.8 | 28.6 ± 17.7 |
| | | BASE | 8.00 | -0.20 ± 0.07 | 205 | 93 | 90.3 ± 1.5 | 49.6 ± 27.6 | 1.2 ± 5.7 | 5.6 ± 6.1 | 38.5 ± 15.1 |
| | | $K = 5$ | 1.80 | -0.20 ± 0.12 | 905 | 643 | 90.0 ± 1.7 | 62.7 ± 13.6 | 8.3 ± 10.7 | 7.0 ± 8.5 | 26.5 ± 17.0 |
| | | $K = L$ | 1.20 | -0.19 ± 0.07 | 935 | 523 | 89.5 ± 1.4 | 58.0 ± 17.2 | 4.9 ± 6.1 | 3.5 ± 3.1 | 29.3 ± 18.8 |
| **CM** | SDO | BASE | 1.30 | -0.52 ± 0.34 | 1018 | 937 | 90.2 ± 1.5 | 63.5 ± 11.5 | 20.3 ± 15.1 | 17.9 ± 14.7 | 26.2 ± 15.8 |
| | | $K = 5$ | 1.15 | -0.53 ± 0.35 | 1020 | 953 | 89.9 ± 1.5 | 63.4 ± 11.5 | 20.4 ± 14.9 | 17.4 ± 14.2 | 26.6 ± 16.0 |
| | | $K = L$ | 1.02 | -0.52 ± 0.35 | 1018 | 951 | 90.2 ± 1.6 | 63.4 ± 12.8 | 18.4 ± 14.1 | 15.6 ± 13.2 | 28.0 ± 16.7 |
| | | BASE | 2.86 | -0.23 ± 0.15 | 757 | 423 | 90.0 ± 1.7 | 58.1 ± 20.5 | 5.7 ± 9.8 | 6.8 ± 7.9 | 28.0 ± 16.6 |
| | | $K = 5$ | 1.55 | -0.24 ± 0.16 | 932 | 609 | 90.0 ± 1.5 | 60.6 ± 16.2 | 7.5 ± 10.2 | 6.4 ± 7.9 | 26.6 ± 17.0 |
| | | $K = L$ | 1.10 | -0.23 ± 0.11 | 988 | 653 | 90.2 ± 1.6 | 59.5 ± 16.8 | 6.3 ± 7.0 | 4.7 ± 4.3 | 28.5 ± 18.3 |
| | | BASE | 6.50 | -0.17 ± 0.07 | 154 | 82 | 90.2 ± 2.0 | 36.7 ± 32.9 | 0.9 ± 4.2 | 3.8 ± 3.9 | 37.3 ± 16.6 |
| | | $K = 5$ | 2.00 | -0.16 ± 0.07 | 740 | 296 | 90.1 ± 1.5 | 55.2 ± 21.2 | 3.1 ± 6.9 | 3.3 ± 3.8 | 29.8 ± 18.0 |
| | | $K = L$ | 1.20 | -0.17 ± 0.06 | 847 | 396 | 90.5 ± 1.6 | 58.8 ± 18.2 | 3.4 ± 5.8 | 2.6 ± 2.5 | 31.1 ± 19.2 |
| **PL** | PROGEN3 | BASE | 1.18 | -0.55 ± 0.40 | 1008 | 910 | 128.0 ± 7.5 | 86.4 ± 15.2 | 37.4 ± 26.7 | 34.2 ± 26.6 | 42.3 ± 25.9 |
| | | $K = 5$ | 1.10 | -0.56 ± 0.40 | 1010 | 940 | 126.0 ± 7.2 | 85.8 ± 13.9 | 36.2 ± 24.6 | 33.3 ± 25.0 | 42.2 ± 25.9 |
| | | $K = L$ | 1.02 | -0.56 ± 0.41 | 1015 | 898 | 115.9 ± 7.5 | 84.6 ± 13.0 | 32.1 ± 24.6 | 30.7 ± 24.7 | 48.5 ± 24.4 |
| | | BASE | 1.50 | -0.32 ± 0.25 | 947 | 787 | 128.2 ± 5.5 | 80.4 ± 16.9 | 22.4 ± 21.2 | 21.2 ± 20.6 | 41.4 ± 26.2 |
| | | $K = 5$ | 1.30 | -0.32 ± 0.25 | 975 | 788 | 127.2 ± 6.2 | 82.8 ± 15.9 | 22.1 ± 21.3 | 20.2 ± 20.6 | 40.7 ± 26.5 |
| | | $K = L$ | 1.04 | -0.32 ± 0.24 | 997 | 800 | 117.7 ± 8.3 | 82.6 ± 16.5 | 18.3 ± 19.0 | 17.4 ± 18.4 | 46.4 ± 25.4 |
| | | BASE | 8.00 | -0.12 ± 0.04 | 62 | 27 | 126.4 ± 3.6 | 32.6 ± 25.8 | 0.4 ± 4.3 | 4.0 ± 3.8 | 68.0 ± 20.5 |
| | | $K = 5$ | 2.20 | -0.12 ± 0.08 | 625 | 312 | 128.5 ± 5.2 | 75.0 ± 24.5 | 4.8 ± 11.3 | 5.3 ± 9.5 | 47.1 ± 28.9 |
| | | $K = L$ | 1.15 | -0.13 ± 0.05 | 863 | 376 | 121.1 ± 6.2 | 77.9 ± 23.0 | 4.2 ± 9.0 | 3.4 ± 4.5 | 50.3 ± 29.2 |
| **PL** | SDO | BASE | 1.50 | -0.37 ± 0.29 | 939 | 749 | 127.5 ± 5.4 | 79.1 ± 18.0 | 19.5 ± 18.9 | 19.3 ± 18.8 | 39.9 ± 26.2 |
| | | $K = 5$ | 1.30 | -0.35 ± 0.30 | 946 | 750 | 127.6 ± 5.5 | 80.9 ± 17.3 | 19.4 ± 19.5 | 18.2 ± 18.7 | 40.4 ± 26.1 |
| | | $K = L$ | 1.02 | -0.34 ± 0.34 | 985 | 720 | 129.9 ± 6.9 | 81.9 ± 24.0 | 16.5 ± 20.2 | 14.9 ± 18.2 | 44.1 ± 27.3 |
| | | BASE | 3.00 | -0.16 ± 0.12 | 526 | 297 | 127.9 ± 4.5 | 61.2 ± 27.9 | 4.9 ± 11.1 | 7.1 ± 10.6 | 48.9 ± 25.3 |
| | | $K = 5$ | 1.80 | -0.16 ± 0.15 | 702 | 421 | 128.8 ± 5.5 | 77.7 ± 23.2 | 7.0 ± 12.3 | 7.3 ± 11.1 | 41.9 ± 27.0 |
| | | $K = L$ | 1.05 | -0.16 ± 0.13 | 901 | 465 | 129.4 ± 6.1 | 80.9 ± 24.1 | 6.2 ± 10.6 | 5.2 ± 7.7 | 46.4 ± 29.4 |
| | | BASE | 5.00 | -0.12 ± 0.61 | 213 | 96 | 126.8 ± 3.7 | 41.4 ± 29.3 | 1.1 ± 5.0 | 3.9 ± 5.7 | 63.9 ± 23.1 |
| | | $K = 5$ | 2.30 | -0.12 ± 0.10 | 500 | 248 | 129.6 ± 5.7 | 74.5 ± 28.7 | 4.1 ± 11.0 | 4.4 ± 8.0 | 44.8 ± 27.2 |
| | | $K = L$ | 1.10 | -0.12 ± 0.07 | 837 | 308 | 129.6 ± 6.3 | 79.5 ± 26.7 | 3.4 ± 7.6 | 2.9 ± 3.9 | 48.6 ± 29.5 |

*Table 4.* **Top-p sampling analysis.** We compare top-$p$ decoding settings for the **sDO** architecture across the CM and PL families. Statistics are computed from 1024 generated sequences for each setting. Reported metrics include the target $\log P$ used for sampling, enabling comparison with the values in Table 3. At matched generative quality, these metrics are very similar to those obtained with standard autoregressive low-temperature decoding.

| Family | $p$ | $\beta$ | Target $\log P$ | Unique (on 1024) | Unique (radius 5) | Length | Dist (Gen–Gen) | Min Dist (Gen–Gen) | Min Dist (Gen–Nat) |
|---|---|---|---|---|---|---|---|---|---|
| **CM** | 1.00 | 2.60 | -0.23 | 744 | 420 | $89.9 \pm 1.7$ | $56.7 \pm 21.7$ | $5.4 \pm 9.1$ | $6.8 \pm 7.4$ |
| | 0.95 | 2.60 | | 727 | 412 | $89.9 \pm 1.7$ | $58.3 \pm 20.5$ | $5.2 \pm 8.9$ | $6.9 \pm 7.5$ |
| | 0.90 | 2.30 | | 707 | 411 | $89.9 \pm 1.6$ | $58.9 \pm 19.8$ | $5.0 \pm 8.8$ | $7.0 \pm 7.7$ |
| | 0.85 | 2.05 | | 735 | 419 | $89.9 \pm 1.6$ | $59.6 \pm 18.8$ | $5.0 \pm 8.3$ | $7.1 \pm 7.6$ |
| | 0.80 | 1.85 | | 735 | 405 | $89.9 \pm 1.5$ | $59.7 \pm 18.9$ | $4.6 \pm 7.9$ | $7.0 \pm 7.4$ |
| **PL** | 1.00 | 3.00 | -0.16 | 527 | 294 | $123.5 \pm 4.3$ | $59.6 \pm 25.9$ | $4.2 \pm 9.5$ | $11.2 \pm 9.2$ |
| | 0.95 | 2.50 | | 557 | 297 | $123.9 \pm 4.4$ | $62.1 \pm 26.2$ | $3.8 \pm 8.5$ | $11.2 \pm 8.9$ |
| | 0.90 | 2.20 | | 563 | 309 | $124.1 \pm 4.6$ | $64.0 \pm 26.1$ | $3.7 \pm 8.3$ | $11.2 \pm 8.7$ |
| | 0.85 | 1.95 | | 540 | 303 | $124.2 \pm 4.8$ | $65.2 \pm 26.9$ | $3.9 \pm 8.2$ | $11.3 \pm 9.4$ |
| | 0.80 | 1.75 | | 530 | 307 | $124.3 \pm 4.8$ | $66.4 \pm 26.6$ | $3.8 \pm 8.4$ | $11.1 \pm 9.0$ |

*Table 5.* **Steering generation toward a Target wild-type (WT).** We report the mean Edit Distance to the target sequence and the generative quality ($\log P$). Statistics are computed over 1024 sequences per sample. Base refers to standard autoregressive sampling, while steered samples are generated using ILMC with parameters $B = 2$, $K = 5$, and $N_{\mathrm{MC}} = 10$.

| Family | Model | Strategy | Dist (to WT) | $\log P$ |
|---|---|---|---|---|
| CM | sDO | Target WT | 0 | -1.19 |
| | | Base ($\beta = 1$) | $64.2 \pm 4.0$ | $-0.83 \pm 0.47$ |
| | | $\beta = 1, \lambda^{\mathrm{WT}} = 2.5$ | $35.5 \pm 15.9$ | $-1.05 \pm 0.34$ |
| | | $\beta = 1, \lambda^{\mathrm{WT}} = 7.5$ | $4.3 \pm 3.7$ | $-1.16 \pm 0.04$ |
| | ProGen3 | Target WT | 0 | -1.08 |
| | | Base ($\beta = 1.1$) | $64.2 \pm 4.0$ | $-0.66 \pm 0.37$ |
| | | $\beta = 1.1, \lambda^{\mathrm{WT}} = 2.5$ | $45.2 \pm 14.0$ | $-0.89 \pm 0.37$ |
| | | $\beta = 1.1, \lambda^{\mathrm{WT}} = 7.5$ | $7.7 \pm 5.6$ | $-1.14 \pm 0.10$ |
| PL | sDO | Target WT | 0 | -1.28 |
| | | Base ($\beta = 1$) | $95.1 \pm 5.9$ | $-0.97 \pm 0.63$ |
| | | $\beta = 1, \lambda^{\mathrm{WT}} = 2.5$ | $74.5 \pm 14.5$ | $-1.22 \pm 0.51$ |
| | | $\beta = 1, \lambda^{\mathrm{WT}} = 7.0$ | $21.9 \pm 12.5$ | $-1.63 \pm 0.29$ |
| | ProGen3 | Target WT | 0 | -1.03 |
| | | Base ($\beta = 1.1$) | $94.5 \pm 7.1$ | $-0.65 \pm 0.44$ |
| | | $\beta = 1.1, \lambda^{\mathrm{WT}} = 4.5$ | $52.9 \pm 21.9$ | $-1.47 \pm 0.35$ |
| | | $\beta = 1.1, \lambda^{\mathrm{WT}} = 10.0$ | $35.5 \pm 21.3$ | $-1.67 \pm 0.30$ |

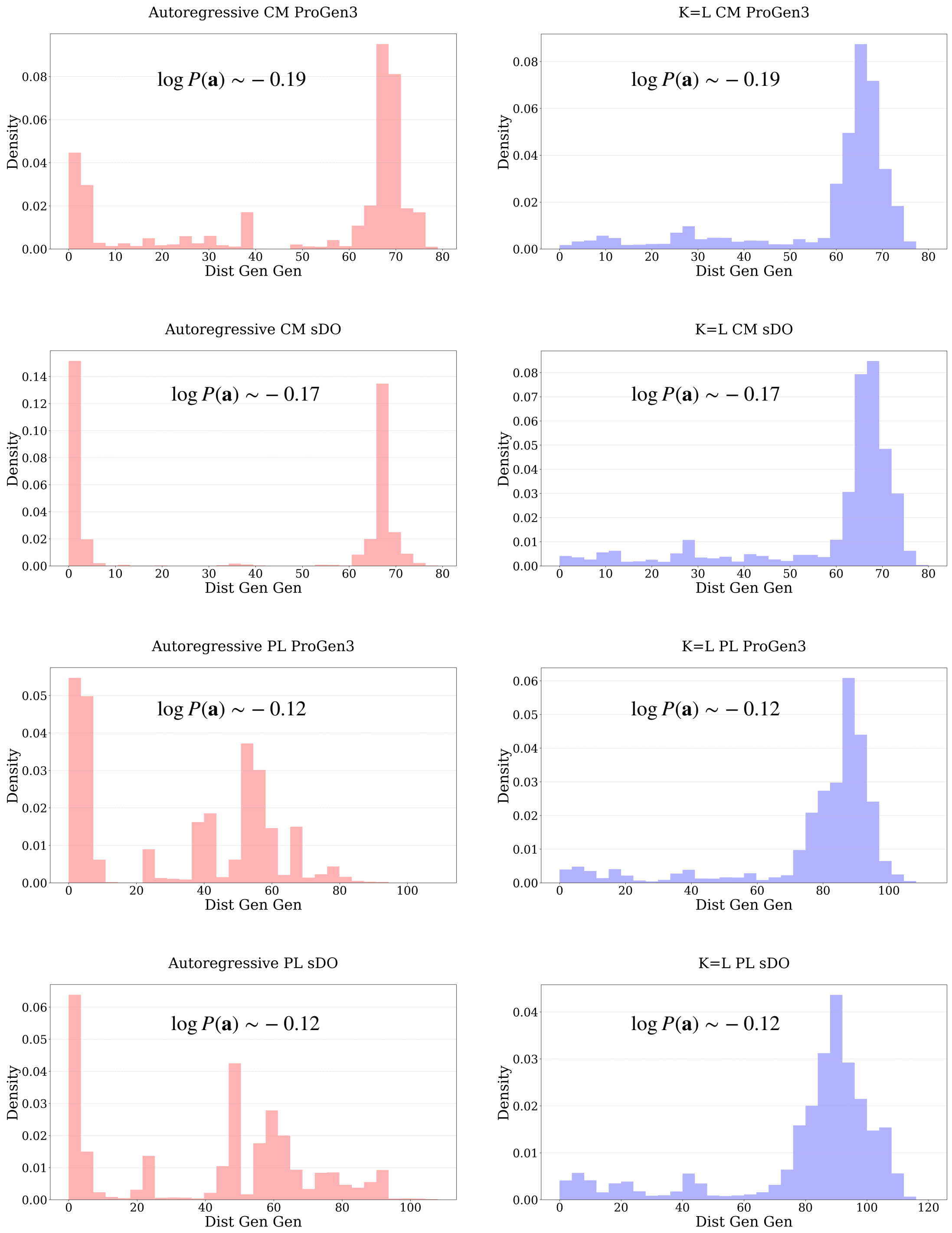


*Figure 6.* **Library intra-distance (Dist Gen-Gen) of the highest** $\log P$ **energy samples from Table 3.** Each row compares standard autoregressive sampling (left, red) and ILMC with $K = L$ ($B = 2$, $N_{\text{MC}} = 10$) (right, blue), matched for the same mean $\log P$. The rows correspond to: (1) ProGen3 on CM, (2) sDO on CM, (3) ProGen3 on PL, and (4) sDO on PL. All samples consist of 1024 sequences. It is evident that the autoregressive sampling (left, red) is collapsing in diversity, as indicated by the consistent peak near 0 pairwise edit distance. ILMC samples (right column) do not exhibit this peak, consistently sampling libraries with typical pairwise distances close to the average distance between natural sequences (91 for PL and 64 for CM).

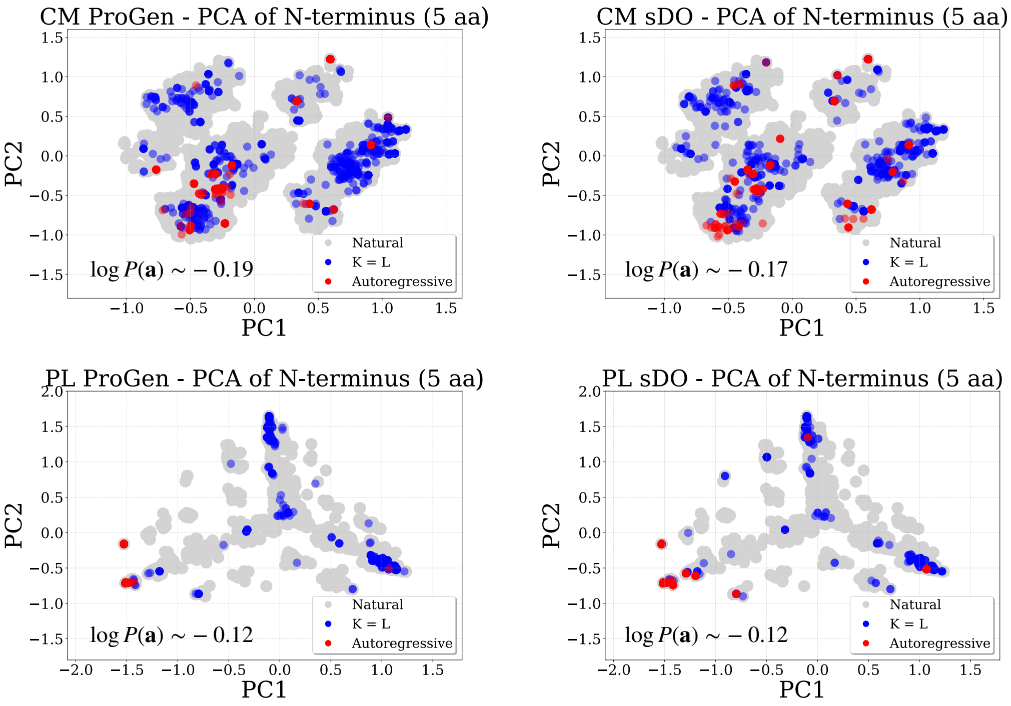


*Figure 7.* **PCA projection of the first 5 amino acids.** The panels correspond to the two families (CM top, PL bottom) and the two models (ProGen3 top, sDO bottom). The projection of the natural data is shown in gray. High $\log P$ samples generated via standard autoregressive sampling are shown in red, while samples generated via ILMC ($B = 2, K = L, N_{\mathrm{MC}} = 10$) are shown in blue. All samples consist of 1024 sequences. These distributions correspond to the highest $\log P$ samples reported in Table 3.

### B. Proofs

#### B.1. Proof of Proposition 2.1

We seek to maximize the entropy $H[Q] = -\sum_{\mathbf{a}} Q(\mathbf{a}) \log Q(\mathbf{a})$ subject to the constraints $\mathbb{E}_Q[E(\mathbf{a})] = C_{\mathrm{E}}$, $\mathbb{E}_Q[S(\mathbf{a})] = C_{\mathrm{S}}$, and the normalization constraint $\sum_{\mathbf{a}} Q(\mathbf{a}) = 1$. The Lagrangian for this optimization problem is:

$$\mathcal{L} = -\sum_{\mathbf{a}} Q(\mathbf{a}) \log Q(\mathbf{a}) - \beta \left( \sum_{\mathbf{a}} Q(\mathbf{a}) E(\mathbf{a}) - C_{\mathrm{E}} \right) - \lambda \left( \sum_{\mathbf{a}} Q(\mathbf{a}) S(\mathbf{a}) - C_{\mathrm{S}} \right) - \gamma \left( \sum_{\mathbf{a}} Q(\mathbf{a}) - 1 \right) \tag{16}$$

Taking the functional derivative with respect to $Q(\mathbf{a})$ and setting it to zero:

$$\frac{\partial \mathcal{L}}{\partial Q(\mathbf{a})} = -\log Q(\mathbf{a}) - 1 - \beta E(\mathbf{a}) - \lambda S(\mathbf{a}) - \gamma = 0 \tag{17}$$

Solving for $Q(\mathbf{a})$ yields:

$$Q^*(\mathbf{a}) = \exp(-1-\gamma) \exp(-\beta E(\mathbf{a}) - \lambda S(\mathbf{a})) \tag{18}$$

By defining the partition function $Z^*(\beta, \lambda) = \exp(1+\gamma)$, so that (18) is a normalized distribution, we recover the form of the canonical Gibbs distribution:

$$Q^*(\mathbf{a}) = \frac{1}{Z^*(\beta, \lambda)} \exp(-\beta E(\mathbf{a}) - \lambda S(\mathbf{a})) \tag{19}$$

#### B.2. Proof of Proposition 2.2

Consider any distribution $Q(\mathbf{a})$ satisfying the constraints $\mathbb{E}_Q[E(\mathbf{a})] = C_{\mathrm{E}}$ and $\mathbb{E}_Q[S(\mathbf{a})] = C_{\mathrm{S}}$. The KL divergence from $Q$ to the optimal $Q^*$ is:

$$\begin{aligned} D_{\mathrm{KL}}(Q \parallel Q^*) &= \sum_{\mathbf{a}} Q(\mathbf{a}) \log \frac{Q(\mathbf{a})}{Q^*(\mathbf{a})} \\ &= -H[Q] - \sum_{\mathbf{a}} Q(\mathbf{a}) \log Q^*(\mathbf{a}) \end{aligned} \tag{20}$$

Substituting the form $\log Q^*(\mathbf{a}) = -\beta E(\mathbf{a}) - \lambda S(\mathbf{a}) - \log Z^*(\beta, \lambda)$:

$$\begin{aligned} D_{\mathrm{KL}}(Q \parallel Q^*) &= -H[Q] - \sum_{\mathbf{a}} Q(\mathbf{a}) \left(-\beta E(\mathbf{a}) - \lambda S(\mathbf{a}) - \log Z^*(\beta, \lambda)\right) \\ &= -H[Q] + \beta \left( \sum_{\mathbf{a}} Q(\mathbf{a}) E(\mathbf{a}) \right) + \lambda \left( \sum_{\mathbf{a}} Q(\mathbf{a}) S(\mathbf{a}) \right) + \log Z^*(\beta, \lambda) \\ &= -H[Q] + \beta C_{\mathrm{E}} + \lambda C_{\mathrm{S}} + \log Z^*(\beta, \lambda) \end{aligned} \tag{21}$$

Since the terms $\beta C_{\mathrm{E}} + \lambda C_{\mathrm{S}} + \log Z^*(\beta, \lambda)$ are independent of $Q$, we have that $D_{\mathrm{KL}}(Q \parallel Q^*)$ is minimized as a function of $Q$ if and only if $H[Q]$ is maximized.

### C. Greedy low-temperature sampling in autoregressive models

A common strategy employed to sample high probability sequences from autoregressive models is *greedy* "low-temperature" autoregressive sampling. Despite the name, it differs from low-temperature sampling in the statistical physics sense (Karan & Du, 2025), which corresponds to sampling from a exponentiated distribution $Q^*(\mathbf{a}) \propto P(\mathbf{a})^\beta$. Here, we analyze the discrepancy between these two strategies and explain why direct autoregressive sampling from the optimal entropy-constrained distribution $Q^*$ is computationally intractable. In this section we focus on the non-steered case (*i.e.*, $\lambda = 0$ in Eq. (1)) for simplicity. The generalization of the analysis discussion to the steered case is very similar and is given in Appendix F.

Let a protein sequence be denoted by $\mathbf{a} = (a_1, \ldots, a_L)$. The pre-trained autoregressive model provides a probability distribution $P(\mathbf{a}) = \prod_{t=1}^{L} P(a_t | \mathbf{a}_{<t})$. Standard low-temperature sampling sharpens the next-token probabilities at each

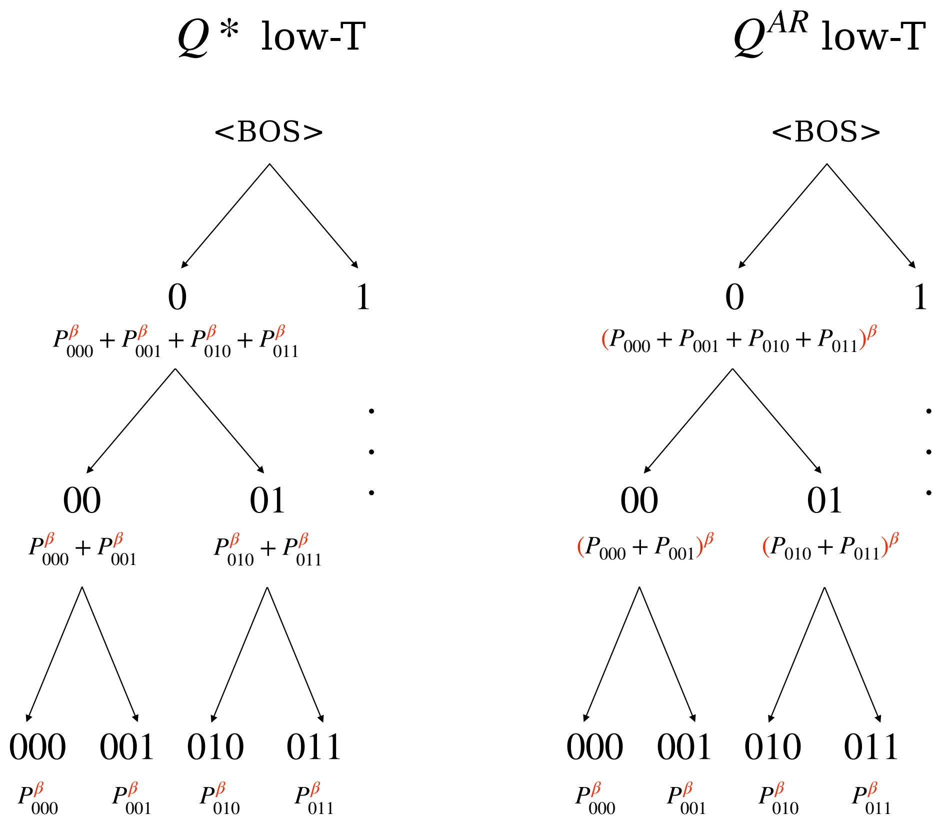


*Figure 8.* Illustrated example of the sampling process as a branching tree decomposition for a binary sequence of length 3. The values at each node represent the relative weight of that branch in relation to the concurrent one. **Left:** The optimal global distribution $Q^*$, derived by exponentiating the probabilities of the leaves. **Right:** The standard autoregressive $Q^{\text{AR}}$, derived by exponentiating local transitions. At any given node, the autoregressive model only provides access to the standard probability mass of its children (the sum in black, corresponding to $\beta = 1$). For the standard strategy (Right), the sampling weights are trivially obtained by exponentiating these local values. However, for the optimal strategy (Left), the correct node weight corresponds to the sum of the exponentiated leaves. Since the summation of probabilities does not commute with exponentiation (i.e., $(\sum p)^\beta \neq \sum p^\beta$), determining the weights for $Q^*$ requires an intractable summation over all future trajectories rather than a simple local operation.

step $t$ toward high-probability tokens by exponentiating their distribution with an inverse temperature parameter $\beta$. Given a prefix $\mathbf{a}_{<t}$, the next token is drawn from a locally re-normalized distribution:

$$Q^{\text{AR}}(a_t|\mathbf{a}_{<t}) = \frac{P(a_t|\mathbf{a}_{<t})^\beta}{Z_t(\mathbf{a}_{<t};\beta)} \tag{22}$$

where $Z_t(\mathbf{a}_{<t};\beta) = \sum_{v\in\mathcal{A}} P(v|\mathbf{a}_{<t})^\beta$ sums only over the vocabulary $\mathcal{A}$. Consequently, the probability of a full sequence generated via this strategy is simply the multiplication of its token draw probabilities:

$$Q^{\text{AR}}(\mathbf{a}) = \prod_{t=1}^{L} \frac{P(a_t|\mathbf{a}_{<t})^\beta}{Z_t(\mathbf{a}_{<t};\beta)} = \frac{P(\mathbf{a})^\beta}{\prod_{t=1}^{L} Z_t(\mathbf{a}_{<t};\beta)} \tag{23}$$

On the other hand, as derived in Proposition 2.1, the unique distribution that maximizes entropy for a fixed mean energy is the Boltzmann distribution $Q^*(\mathbf{a}) = P(\mathbf{a})^\beta / Z^*(\beta)$, where $Z^*(\beta)$ is the global partition function summing over all possible sequences in the space $\mathcal{A}^L$.

To understand why $Q^{\text{AR}} \neq Q^*$, we must examine the conditional probability of the next token $a_t$ from the optimal distribution $Q^*$. By definition, $Q^*(a_t|\mathbf{a}_{<t}) = Q^*(\mathbf{a}_{\leq t})/Q^*(\mathbf{a}_{<t})$. Here, the marginal probability of a prefix is obtained by

summing the global probabilities of all completed sequences (the ”leaves”) rooted at that prefix. We derive the expression for the optimal conditional probability starting from this definition:

$$Q^*(a_t|\mathbf{a}_{<t}) = \frac{Q^*(\mathbf{a}_{\leq t})}{Q^*(\mathbf{a}_{<t})} \tag{24}$$

$$= \frac{\sum_{\mathbf{a}'_{>t}} Q^*(\mathbf{a}_{\leq t}, \mathbf{a}'_{>t})}{\sum_{a'_t} \sum_{\mathbf{a}'_{>t}} Q^*(\mathbf{a}_{<t}, a'_t, \mathbf{a}'_{>t})} \tag{25}$$

Substituting the definition of the global Boltzmann distribution $Q^*(\mathbf{a}) = \frac{1}{Z^*(\beta)} P(\mathbf{a})^\beta$:

$$Q^*(a_t|\mathbf{a}_{<t}) = \frac{\frac{1}{Z^*(\beta)} \sum_{\mathbf{a}'_{>t}} P(\mathbf{a}_{\leq t}, \mathbf{a}'_{>t})^\beta}{\frac{1}{Z^*(\beta)} \sum_{a'_t} \sum_{\mathbf{a}'_{>t}} P(\mathbf{a}_{<t}, a'_t, \mathbf{a}'_{>t})^\beta} \tag{26}$$

$$= \frac{\sum_{\mathbf{a}'_{>t}} \left(P(\mathbf{a}_{\leq t}) P(\mathbf{a}'_{>t}|\mathbf{a}_{\leq t})\right)^\beta}{\sum_{a'_t} \sum_{\mathbf{a}'_{>t}} \left(P(\mathbf{a}_{<t}, a'_t) P(\mathbf{a}'_{>t}|\mathbf{a}_{<t}, a'_t)\right)^\beta} \tag{27}$$

We factor out the prefix probabilities which are constant with respect to the summation over future trajectories $\mathbf{a}'_{>t}$:

$$Q^*(a_t|\mathbf{a}_{<t}) = \frac{P(\mathbf{a}_{\leq t})^\beta \sum_{\mathbf{a}'_{>t}} P(\mathbf{a}'_{>t}|\mathbf{a}_{\leq t})^\beta}{\sum_{a'_t} \left[P(\mathbf{a}_{<t}, a'_t)^\beta \sum_{\mathbf{a}'_{>t}} P(\mathbf{a}'_{>t}|\mathbf{a}_{<t}, a'_t)^\beta\right]} \tag{28}$$

The denominator is a normalization constant involving a summation over all possible next tokens $a'_t$. To find the proportionality relation for a specific $a_t$, we focus on the numerator. Using the decomposition $P(\mathbf{a}_{\leq t}) = P(\mathbf{a}_{<t}) P(a_t|\mathbf{a}_{<t})$:

$$Q^*(a_t|\mathbf{a}_{<t}) \propto P(\mathbf{a}_{\leq t})^\beta \sum_{\mathbf{a}'_{>t}} P(\mathbf{a}'_{>t}|\mathbf{a}_{\leq t})^\beta \tag{29}$$

$$\propto \left(P(\mathbf{a}_{<t}) P(a_t|\mathbf{a}_{<t})\right)^\beta \sum_{\mathbf{a}'_{>t}} P(\mathbf{a}'_{>t}|\mathbf{a}_{\leq t})^\beta \tag{30}$$

Since $P(\mathbf{a}_{<t})^\beta$ depends only on the fixed history and is constant with respect to the choice of $a_t$, it can be absorbed into the proportionality constant. Defining the lookahead term,

$$\mathcal{Z}_{\text{future}}(a_t, \mathbf{a}_{<t}) \equiv \sum_{\mathbf{a}'_{>t}} P(\mathbf{a}'_{>t}|\mathbf{a}_{\leq t})^\beta, \tag{31}$$

we arrive at the final relation:

$$Q^*(a_t|\mathbf{a}_{<t}) \propto \underbrace{P(a_t|\mathbf{a}_{<t})^\beta}_{\text{Autoregressive Term}} \times \underbrace{\mathcal{Z}_t^{\text{future}}(a_t, \mathbf{a}_{<t})}_{\text{Lookahead Term}} \tag{32}$$

Equation (32) highlights the fundamental flaw in greedy autoregressive low-temperature sampling. $Q^{\text{AR}}$ selects the next token based solely on the Autoregressive Term, effectively assuming that $\mathcal{Z}_{\text{future}}(a_t, \mathbf{a}_{<t})$ does not depend on $a_t$. However, $\mathcal{Z}_t^{\text{future}}(a_t, \mathbf{a}_{<t})$ may vary significantly: it measures the “volume” of high-probability paths accessible from the current state, which in some cases may depend critically on the choice of the current token $a_t$. Autoregressive sampling is therefore myopic; it sharpens the next token distribution based only on the immediate edge in the probability tree (Figure 8, right) and may select a token that has high local probability but leads to a ”dead end” in the energy landscape. In contrast, global sampling is prescient, re-weighting the immediate choice $a_t$ by the exponentiated weight of the reachable ending sequences (leaves).

To illustrate this effect, consider a simple scenario where, conditioned on a specific prefix, there are $m$ equally probable future completions (paths). The future weight term behaves as:

$$\mathcal{Z}^{\text{future}} \approx \sum_{i=1}^{m} \left(\frac{1}{m}\right)^\beta = m \cdot m^{-\beta} = m^{1-\beta} \tag{33}$$

Since $\beta > 1$, the exponent $1 - \beta$ is negative. This implies that as $m$ increases (representing a more diffuse or uncertain future), the term $\mathcal{Z}_{\text{future}}$ decreases. Consequently, the optimal distribution $Q^*$ penalizes paths that lead to high-entropy futures, favoring sequences that converge toward specific, high-likelihood outcomes. The greedy autoregressive sampler $Q^{\text{AR}}$ is blind to this downstream uncertainty.

This creates an apparent paradox: $Q^*$ is the unique maximum entropy distribution for a fixed mean generative quality, yet it penalizes diffuse (high-entropy) futures more aggressively than $Q^{\text{AR}}$. The resolution lies in the comparison of the sampling parameters. To achieve an identical mean generative energy using the myopic autoregressive strategy $Q^{\text{AR}}$, one must employ a significantly higher inverse temperature $\beta_{AR} \gg \beta$. This excessive local sharpening in $Q^{\text{AR}}$ collapses the distribution's support early in the generation process, severely restricting diversity.

This analysis also explains why sampling from $Q^*$ in a single autoregressive pass is impossible. Evaluating $\mathcal{Z}_t^{\text{future}}(\mathbf{a}_{\leq t})$ requires summing over $|\mathcal{A}|^{L-t}$ future trajectories (see Figure 8), which is computationally intractable for typical protein sequence lengths. This justifies the need for the ILMC method, which approximates this global distribution.

We also note that $\frac{1}{1-\beta} \log \mathcal{Z}_t^{\text{future}}$ is mathematically equivalent to the Rènyi entropy of the distribution $P(\mathbf{a}_{>t}|\mathbf{a}_{\leq t})$ of continuations $\mathbf{a}_{>t}$ conditioned on a given prefix $\mathbf{a}_{\leq t}$. Rènyi entropies are often used as diversity measures (Hill, 1973), in part because of the following well-known properties:

- For $\beta > 1$, the Rènyi entropy is larger for prefixes $\mathbf{a}_{\leq t}$ for which the conditional distribution $P(\mathbf{a}_{>t}|\mathbf{a}_{\leq t})$ has few but high-probability continuations $\mathbf{a}_{>t}$.
- For $\beta < 1$, the Rènyi entropy is larger for prefixes $\mathbf{a}_{\leq t}$ for which the conditional distribution $P(\mathbf{a}_{>t}|\mathbf{a}_{\leq t})$ has many but low-probability continuations $\mathbf{a}_{>t}$.
- As $\beta \to 1$, the Rènyi entropy approaches the Shannon entropy, which is maximized when the conditional distribution $P(\mathbf{a}_{>t}|\mathbf{a}_{\leq t})$ becomes a flat uniform measure over the possible continuations $\mathbf{a}_{>t}$.

The behavior of $\mathcal{Z}_t^{\text{future}}$ at $\beta > 1$ drives $Q^*$ towards high-probability sequences.

### D. Is $K > 1$ Always Closer to the Optimal Distribution?

Based on Proposition 2.2, minimizing the KL divergence ($D_{\text{KL}}$) between a sampling strategy and the optimal distribution $Q^*$ is equivalent to maximizing the entropy of the sampled library for a fixed generative energy. Since the ILMC method samples from $Q^*$ in the limit where the lookback window $K$ equals the sequence length $L$ and the number of MC steps $N_{\text{MC}} \to \infty$, it is intuitive to hypothesize that increasing $K$ monotonically improves the approximation of $Q^*$. Specifically, one might expect that a strategy with a larger lookback window $K$ would always yield higher entropy for a given target energy compared to a more myopic strategy.

While this intuition holds true in our computational experiments, as shown in Section 3, it is not universally correct. We demonstrate this with a counter-example where $K = 2$ and $B = 2$ (effectively corresponding to an autoregressive strategy that samples token pairs rather than single tokens) yields lower entropy with respect to the standard autoregressive strategy for a given level of mean energy. Consider a binary sequence space of length $L = 4$ governed by the following probability distribution $P(\mathbf{a})$:

$$P(\mathbf{a}) = \begin{cases} 0100 & \text{with prob } 0.44 \quad \text{(Ground State A)} \\ 0110 & \text{with prob } 0.06 \\ 1010 & \text{with prob } 0.44 \quad \text{(Ground State B)} \\ 1100 & \text{with prob } 0.06 \\ \text{others} & \text{with prob } 0.00 \end{cases} \tag{34}$$

Let's consider the limit where the inverse temperature goes to infinity $\beta \to \infty$. In this situation the sampler is forced to strictly follow the path of highest total probability mass. We observe a critical divergence in behavior: the single-token sampler preserves the symmetry between the two ground states, whereas the block sampler breaks this symmetry, leading to mode collapse.

**Case 1: Single-Token Autoregressive Sampling.** The decision process for the single-token sampler ($B = 1$) starts on the first token. The marginal probability of starting with `0` is the sum of probabilities for sequences `0100` and `0110` ($0.44 + 0.06 = 0.50$). Similarly, the marginal probability of starting with `1` is the sum for `1010` and `1100` ($0.44 + 0.06 = 0.50$). Since the initial masses are identical ($0.50$ vs $0.50$), the sampler selects the initial token with equal probability. Consequently, the sampler can settle into either Ground State A or Ground State B with equal likelihood ($P = 0.5$). The resulting entropy is $\log 2$. See Figure 9.

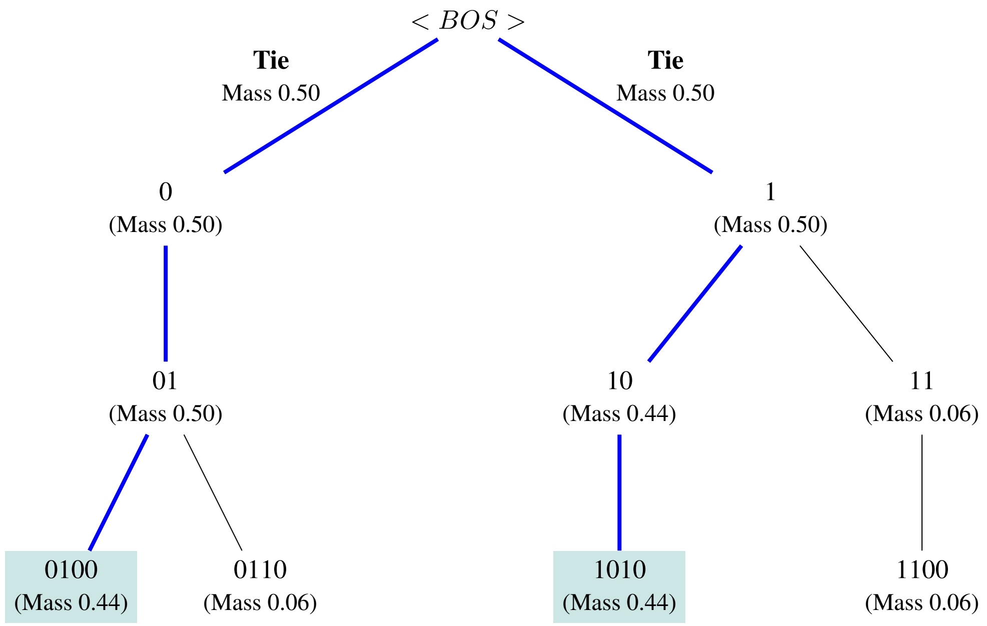


*Figure 9.* Case 1: Single-Token Autoregressive Sampling.

**Case 2: $K = 2$, $B = 2$ Sampling.** Now, consider a block size of $B = 2$ tokens. The sampler compares the joint probabilities of the prefixes $(x_1, x_2)$ directly. The valid prefixes are `01`, `10`, and `11`.

Here, the symmetry is broken. The block `01` aggregates the mass of the dominant state A and the minor state ($0.44 + 0.06 = 0.50$). However, due to the sequence structure, the path to Ground State B requires the block `10`, which has a mass of only $0.44$.

Because $0.50 > 0.44$, the block sampler in the limit of high $\beta$ will exclusively select the `01` branch. It essentially "blindly" discards Ground State B because its associated prefix mass is lower, even though the final state probability is identical to Ground State A. The entropy for this strategy drops to $0$. See Figure 10.

This phenomenon is not limited to the ground state limit but persists across intermediate energy regimes. Figure 11 demonstrates that the single-token strategy yields strictly higher entropy for a given energy level compared to the block-wise approach.

### E. Good and Bad Steering Potentials: Illustrative Examples

Here we illustrate two example steering potentials, a *bad* one that results in a poor sampling strategy, and a *good* one that guides the sampling effectively.

If the steering potential only depends on the terminal portion of the sequence, the early stages of generation will lack guidance, potentially driving the chain into regions of the sequence space where the desired characteristic is no longer reachable. To illustrate this necessity, consider a situation in which we want to steer the generation toward sequences that exhibit a specific target motif, defined as a fixed reference sub-sequence $\mathbf{s} = (s_m, \ldots, s_l)$ residing within the position

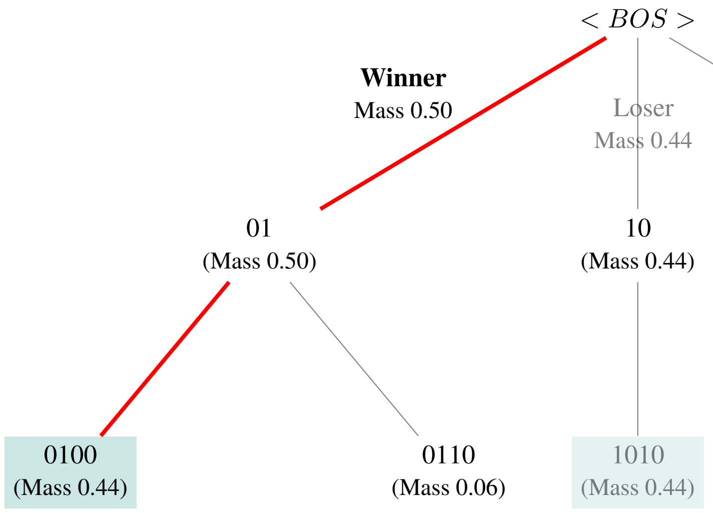


*Figure 10.* Case 2: $K = 2$, $B = 2$ sampling.

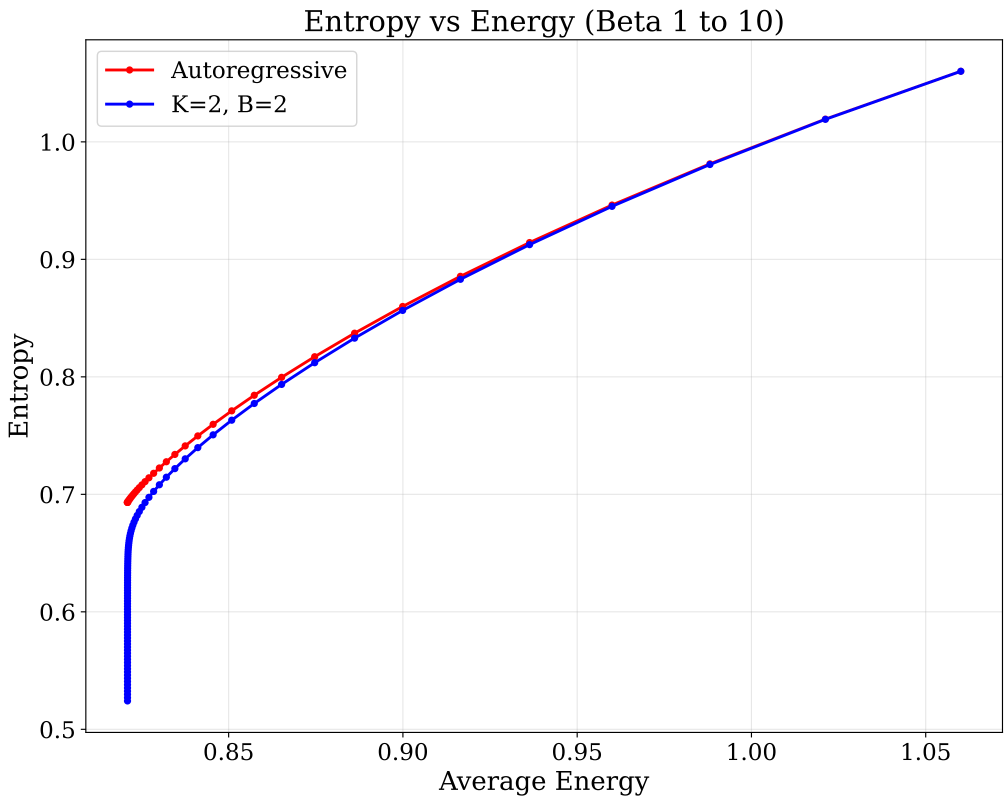


*Figure 11.* Comparison of the Energy-Entropy frontier between the standard single-token autoregressive sampler and the block-wise Iterative Lookback sampler ($K = 2, B = 2$) using as baseline the example distribution $P(a)$. As the inverse temperature $\beta$ increases from 1 to 10 in increments of 0.1, the autoregressive sampler maintains higher entropy at given average energy values compared to the $K = 2, B = 2$ ILMC strategy.

window $[m, l]$. A natural steering potential to quantify the deviation from this target is the Hamming distance:

$$S^{\text{bad}}(\mathbf{a}) = \sum_{i=m}^{l} \mathbb{I}(a_i \neq s_i) \tag{35}$$

The corresponding partial steering potential $S_t^{\text{bad}}(\mathbf{a}_{\leq t})$ can be defined as the Hamming distance to the motif evaluated only on the residues currently generated. For any $t < m$, the partial potential $S_t^{\text{bad}}(\mathbf{a}_{\leq t})$ is trivially constant. Consequently, the refinement steps during the first $m$ tokens provide no steering. The model generates a prefix guided solely by the base generative energy $E_t(\mathbf{a}_{\leq t})$. By the time the sampler reaches index $m$, the prefix may be structurally incompatible with the required motif, leading to a poor generative quality.

To illustrate an example of an effective steering potential, consider the case where the objective is to modify the amino acid composition of the generated proteins, for instance, by increasing the frequency of a desired amino acid $A$. The global score $S^{\text{good}}(\mathbf{a})$ is the negative total count of $A$, while the partial score is the cumulative count:

$$S_t^{\text{good}}(\mathbf{a}_{\leq t}) = -\sum_{i=1}^{t} \mathbb{I}(a_i = A) \tag{36}$$

In this case, the steering is effective from the very first token. The refinement steps immediately bias the sequence toward a higher frequency of $A$, ensuring that the early prefix correlates with the desired final outcome.

### F. What makes a steering potential effective?

Practically, steering potentials are most effective when their prefix form provides an informative signal throughout generation, so that early token choices do not create large, hard-to-correct differences in the remaining achievable score. In this section we attempt to make this intuition more precise. In doing so we generalize some of the calculations from Section C, which did not consider steering, to the steered case.

We begin by recalling the target distribution, which we write as:

$$Q^*(\mathbf{a}) = \frac{1}{Z^*} e^{-U(\mathbf{a})}, \quad \text{with} \quad U(\mathbf{a}) = \beta E(\mathbf{a}) + \lambda S(\mathbf{a}), \tag{37}$$

as introduced in (1), where

$$Z^* = \sum_{\mathbf{a}} e^{-U(\mathbf{a})} \tag{38}$$

is a normalization constant. We suppose that we can decompose these potentials as follows:

$$E(\mathbf{a}) = \sum_i e_i(\mathbf{a}_{\leq i}), \quad S(\mathbf{a}) = \sum_i s_i(\mathbf{a}_{\leq i}), \quad U(\mathbf{a}) = \sum_i u_i(\mathbf{a}_{\leq i}) \tag{39}$$

where the sum considers the contribution of each token, conditioned on the preceding tokens. Note that such a decomposition is formally always possible. For example, given an arbitrary potential $S(\mathbf{a})$, it suffices to define

$$s_i(\mathbf{a}_{\leq i}) = \begin{cases} 0 & i < L \\ S(\mathbf{a}) & i = L \end{cases} \tag{40}$$

Then Eq. (39) is trivially satisfied. But although this definition is mathematically compatible with (39), it would be terrible from the point of view of the ILMC algorithm we introduce in this work, as we will see.

On the other hand, there are several ways to achieve a decomposition like (39). For example, given arbitrary functions $\phi_0, \ldots, \phi_L$, we have that

$$u_i'(\mathbf{a}_{\leq i}) = u_i(\mathbf{a}_{\leq i}) + \phi_i(\mathbf{a}_{\leq i}) - \phi_{i-1}(\mathbf{a}_{\leq i-1}) \tag{41}$$

is also compatible with the global potential when summed across the full sequence,

$$\sum_i u_i'(\mathbf{a}_{\leq i}) = \sum_i u_i(\mathbf{a}_{\leq i}) = U(\mathbf{a}) \tag{42}$$

provided that $\phi_0 \equiv 0$ and $\phi_L \equiv 0$. This telescoping gauge invariance may allow us to reallocate lookahead information into prefix terms, which in turn makes ILMC more effective than a naive decomposition such as Eq. (40).

As in the main-text, we consider a generated prefix $\mathbf{a}_{\leq t}$, and define the partial quantities:

$$E_t(\mathbf{a}_{\leq t}) = \sum_{i \leq t} e_i(\mathbf{a}_{\leq i}), \quad S_t(\mathbf{a}_{\leq t}) = \sum_{i \leq t} s_i(\mathbf{a}_{\leq i}), \quad U_t(\mathbf{a}_{\leq t}) = \sum_{i \leq t} u_i(\mathbf{a}_{\leq i}) \tag{43}$$

Now, substituting the decomposition (39), the target distribution $Q^*$ can be written:

$$Q^*(\mathbf{a}) = \frac{1}{Z^*} \exp\left\{ -\sum_j u_j(\mathbf{a}_{\leq j}) \right\} \tag{44}$$

We focus on the generation of a token at site $i$. To emphasize the dependence on $a_i$, we split the exponent as follows:

$$Q^*(\mathbf{a}) = \frac{1}{Z^*} \exp\left\{ -\sum_{j<i} u_j(\mathbf{a}_{\leq j}) - u_i(a_i, \mathbf{a}_{<i}) - \sum_{j>i} u_j(\mathbf{a}_{i+1:j}, a_i, \mathbf{a}_{<i}) \right\} \tag{45}$$

Consider the next-token distribution, $Q_i^*(a_i|\mathbf{a}_{<i})$. Following calculations very similar to those already given in Sec. C,

$$\begin{aligned}
Q_i^*(a_i|\mathbf{a}_{<i}) &= \frac{\sum_{\mathbf{a}'_{>i}} Q^*(\mathbf{a}'_{>i}, a_i, \mathbf{a}_{<i})}{\sum_{\mathbf{a}'_{\geq i}} Q^*(\mathbf{a}'_{>i}, a'_i, \mathbf{a}_{<i})} &(46)\\
&= \frac{\sum_{\mathbf{a}'_{>i}} \exp\left\{ -\sum_{j<i} u_j(\mathbf{a}_{\leq j}) - u_i(a_i, \mathbf{a}_{<i}) - \sum_{j>i} u_j(\mathbf{a}'_{i+1:j}, a_i, \mathbf{a}_{<i}) \right\}}{\sum_{\mathbf{a}'_{\geq i}} \exp\left\{ -\sum_{j<i} u_j(\mathbf{a}_{\leq j}) - u_i(a'_i, \mathbf{a}_{<i}) - \sum_{j>i} u_j(\mathbf{a}'_{i+1:j}, a'_i, \mathbf{a}_{<i}) \right\}} &(47)\\
&= \exp\left\{-u_i(a_i, \mathbf{a}_{<i})\right\} \frac{\sum_{\mathbf{a}'_{>i}} \exp\left\{ -\sum_{j>i} u_j(\mathbf{a}'_{i+1:j}, a_i, \mathbf{a}_{<i}) \right\}}{\sum_{\mathbf{a}'_{\geq i}} \exp\left\{ -u_i(a'_i, \mathbf{a}_{<i}) - \sum_{j>i} u_j(\mathbf{a}'_{i+1:j}, a'_i, \mathbf{a}_{<i}) \right\}} &(48)\\
&= Q_i^{\mathrm{AR}}(a_i|\mathbf{a}_{<i}) \frac{\mathcal{Z}_i^{\mathrm{future}}(a_i, \mathbf{a}_{<i})}{\mathbb{E}_{a'_i \sim Q_i(a'_i|\mathbf{a}_{<i})} \mathcal{Z}_i^{\mathrm{future}}(a'_i, \mathbf{a}_{<i})} &(49)
\end{aligned}$$

where

$$Q_i^{\mathrm{AR}}(a_i|\mathbf{a}_{<i}) = \frac{\exp\left\{-u_i(a_i, \mathbf{a}_{<i})\right\}}{\sum_{a'_i} \exp\left\{-u_i(a'_i, \mathbf{a}_{<i})\right\}} \tag{50}$$

is the greedy autoregressive sampler, and

$$\mathcal{Z}_i^{\mathrm{future}}(\mathbf{a}_{\leq i}) = \sum_{\mathbf{a}_{>i}} \exp\left\{ -\sum_{j>i} u_j(\mathbf{a}_{\leq j}) \right\} \tag{51}$$

Note that in contrast to the previously defined (23) and (31) from Sec. C, we now incorporate the prefix steering potential as part of $Q^{\mathrm{AR}}$.

To keep the calculation simple, we have effectively assumed that autoregressive sampling is able to incorporate the prefix steering potential exactly in the next-token proposal. Note that this can always be achieved by making $N_{\mathrm{MC}}$ sufficiently large, regardless of the block-size $B$ or the lookback window $K$ parameters of ILMC. Even if the next-token proposal incorporates steering exactly, the sampling strategy may still deviate significantly from the target $Q^*$, due to the finiteness of $K$ or $B$. In the calculation that follows, we can either assume that $B = K = 1$, or that the tokens actually represent blocks of size $B = K$.

Thus, now the goal is to compare the autoregressive sampler:

$$Q^{\mathrm{AR}}(\mathbf{a}) = \prod_i Q_i^{\mathrm{AR}}(a_i|\mathbf{a}_{<i}) \tag{52}$$

to the target distribution $Q^*$. To do so, we compute their KL divergence. Since we also have $Q^*(\mathbf{a}) = \prod_i Q_i^*(a_i|\mathbf{a}_{<i})$, we can write:

$$\begin{aligned}
D_{\mathrm{KL}}(Q^{\mathrm{AR}} \| Q^*) &= \sum_{\mathbf{a}} Q^{\mathrm{AR}}(\mathbf{a}) \log \frac{Q^{\mathrm{AR}}(\mathbf{a})}{Q^*(\mathbf{a})} = \sum_i \sum_{\mathbf{a}} Q^{\mathrm{AR}}(\mathbf{a}_{<i}) Q_i^{\mathrm{AR}}(a_i|\mathbf{a}_{<i}) \log \frac{Q_i^{\mathrm{AR}}(a_i|\mathbf{a}_{<i})}{Q_i^*(a_i|\mathbf{a}_{<i})} &(53)\\
&= \sum_i \mathbb{E}_{\mathbf{a}_{<i} \sim Q^{\mathrm{AR}}} D_{\mathrm{KL}}\left( Q_i^{\mathrm{AR}}(\cdot|\mathbf{a}_{<i}) \| Q_i^*(\cdot|\mathbf{a}_{<i}) \right) &(54)
\end{aligned}$$

with

$$D_{\mathrm{KL}}\left(Q_i^{\mathrm{AR}}(\cdot|\mathbf{a}_{<i})\|Q_i^*(\cdot|\mathbf{a}_{<i})\right)=\sum_{a_i}Q_i^{\mathrm{AR}}(a_i|\mathbf{a}_{<i})\log\frac{Q_i^{\mathrm{AR}}(a_i|\mathbf{a}_{<i})}{Q_i^*(a_i|\mathbf{a}_{<i})} \tag{55}$$

$$=-\sum_{a_i}Q_i^{\mathrm{AR}}(a_i|\mathbf{a}_{<i})\log\frac{\mathcal{Z}_i^{\mathrm{future}}(a_i,\mathbf{a}_{<i})}{\mathbb{E}_{a_i'\sim Q_i^{\mathrm{AR}}(a_i'|\mathbf{a}_{<i})}\mathcal{Z}_i^{\mathrm{future}}(a_i',\mathbf{a}_{<i})} \tag{56}$$

$$=\log\mathbb{E}_{a_i\sim Q_i^{\mathrm{AR}}(a_i|\mathbf{a}_{<i})}\mathcal{Z}_i^{\mathrm{future}}(a_i,\mathbf{a}_{<i})-\mathbb{E}_{a_i\sim Q_i^{\mathrm{AR}}(a_i|\mathbf{a}_{<i})}\log\mathcal{Z}_i^{\mathrm{future}}(a_i,\mathbf{a}_{<i}) \tag{57}$$

Therefore, $Q_i^{\mathrm{AR}}(a_i|\mathbf{a}_{<i})$ equals $Q_i^*(a_i|\mathbf{a}_{<i})$ if and only if $\mathcal{Z}_i^{\mathrm{future}}(a_i,\mathbf{a}_{<i})$ does not depend on $a_i$ within the support of $Q_i^{\mathrm{AR}}(a_i|\mathbf{a}_{<i})$. Otherwise, any dependence of the lookahead term $\mathcal{Z}_i^{\mathrm{future}}(a_i,\mathbf{a}_{<i})$ on the current token $a_i$, manifests itself in a larger KL divergence between the autoregressive proposal $Q_i^{\mathrm{AR}}(a_i|\mathbf{a}_{<i})$ and the target $Q_i^*(a_i|\mathbf{a}_{<i})$, which in turn results in global errors in (53).

An easy bound on the error induced by the non-flatness of the lookahead term is obtained by defining:

$$r_i(\mathbf{a}_{<i})=\log\frac{\max_{a_i}\mathcal{Z}_i^{\mathrm{future}}(a_i,\mathbf{a}_{<i})}{\min_{a_i}\mathcal{Z}_i^{\mathrm{future}}(a_i,\mathbf{a}_{<i})} \tag{58}$$

Then it follows from (55) that:

$$D_{\mathrm{KL}}\left(Q_i^{\mathrm{AR}}(\cdot|\mathbf{a}_{<i})\|Q_i^*(\cdot|\mathbf{a}_{<i})\right)\leq\log\max_{a_i}\mathcal{Z}_i^{\mathrm{future}}(a_i,\mathbf{a}_{<i})-\log\min_{a_i}\mathcal{Z}_i^{\mathrm{future}}(a_i,\mathbf{a}_{<i})=r_i(\mathbf{a}_{<i}) \tag{59}$$

If we furthermore have a uniform bound over all prefixes:

$$R_i=\max_{\mathbf{a}_{<i}}r_i(\mathbf{a}_{<i}) \tag{60}$$

then an immediate consequence of (53) is that:

$$D_{\mathrm{KL}}(Q^{\mathrm{AR}}\|Q^*)\leq\sum_i R_i \tag{61}$$

which bounds the global error on the approximation of the target distribution.

Although computing this error bound may be difficult in practice, this calculation proves that making $\mathcal{Z}_i^{\mathrm{future}}(a_i,\mathbf{a}_{<i})$ as flat as possible as a function of $a_i$, or more precisely minimizing (60), brings the distribution $Q^{\mathrm{AR}}$ closer to the target.

We can also obtain a bound that relates more directly to the variation of the potential terms $u_j(\mathbf{a}_{\leq j})$. Defining:

$$u_{i\to j}^{\max}(\mathbf{a}_{<i},\mathbf{a}_{i+1:j})=\max_{a_i}u_j(\mathbf{a}_{<i},a_i,\mathbf{a}_{i+1:j}),\quad u_{i\to j}^{\min}(\mathbf{a}_{<i},\mathbf{a}_{i+1:j})=\min_{a_i}u_j(\mathbf{a}_{<i},a_i,\mathbf{a}_{i+1:j}) \tag{62}$$

for $i<j$, it is clear that:

$$\sum_{\mathbf{a}_{>i}}\exp\left\{-\sum_{j>i}u_{i\to j}^{\max}(\mathbf{a}_{<i},\mathbf{a}_{i+1:j})\right\}\leq\mathcal{Z}_i^{\mathrm{future}}(\mathbf{a}_{\leq i})\leq\sum_{\mathbf{a}_{>i}}\exp\left\{-\sum_{j>i}u_{i\to j}^{\min}(\mathbf{a}_{<i},\mathbf{a}_{i+1:j})\right\} \tag{63}$$

Hence,

$$r_i(\mathbf{a}_{<i})\leq\log\frac{\sum_{\mathbf{a}_{i+1:j}}\exp\left\{-\sum_{j>i}u_{i\to j}^{\min}(\mathbf{a}_{<i},\mathbf{a}_{i+1:j})\right\}}{\sum_{\mathbf{a}_{i+1:j}}\exp\left\{-\sum_{j>i}u_{i\to j}^{\max}(\mathbf{a}_{<i},\mathbf{a}_{i+1:j})\right\}}. \tag{64}$$

Next, defining:

$$\Delta_{i\to j}(\mathbf{a}_{<i})=\max_{\mathbf{a}_{i+1:j}}\left\{u_{i\to j}^{\max}(\mathbf{a}_{<i},\mathbf{a}_{i+1:j})-u_{i\to j}^{\min}(\mathbf{a}_{<i},\mathbf{a}_{i+1:j})\right\} \tag{65}$$

$$=\max_{\mathbf{a}_{i+1:j}}\max_{a_i,a_i'}|u_j(\mathbf{a}_{<i},a_i,\mathbf{a}_{i+1:j})-u_j(\mathbf{a}_{<i},a_i',\mathbf{a}_{i+1:j})| \tag{66}$$

we have that $u_{i\to j}^{\max}(\mathbf{a}_{<i}, \mathbf{a}_{i+1:j}) \le u_{i\to j}^{\min}(\mathbf{a}_{<i}, \mathbf{a}_{i+1:j}) + \Delta_{i\to j}(\mathbf{a}_{<i})$, and therefore,

$$r_i(\mathbf{a}_{<i}) \le \sum_{j>i} \Delta_{i\to j}(\mathbf{a}_{<i}) \tag{67}$$

which gives a bound directly in terms of the "lookahead" variations $\Delta_{i\to j}(\mathbf{a}_{<i})$ in the potentials. In turn, these can be used in Equations (60) and (61) to bound the global KL divergence directly.

As an example, consider a potential of the simple form:

$$U(\mathbf{a}) = \sum_i u_i(a_i) \tag{68}$$

a particular case of Eq. (39) where the contribution to the potential of each token can be made independent of previous tokens. By substituting (68) into (51), one sees immediately that $\mathcal{Z}_i^{\text{future}}(a_i, \mathbf{a}_{<i})$ becomes independent of $a_i$. Indeed, $\Delta_{i\to j}(\mathbf{a}_{<i}) \equiv 0$ in this simple example. The autoregressive strategy then matches the target $Q^*$ perfectly.

However, one could also reorganize the potential decomposition (68) using the telescopic gauge invariance (41), bringing it to the form (40). In this case, all the lookahead terms exhibit dependences on the current token $a_i$, which weakens the bounds (60) and results in a much larger global divergence (53). This example shows that, besides the definition of the potential itself, the way in which the decomposition (39) is constructed also plays a crucial role in the effectiveness of the resulting sampling strategy.

In the framework of ILMC that we have introduced here, the generative potential $E(\mathbf{a})$ and its prefix decomposition are fixed by the autoregressive form of the base generative model. However, we can still reduce dependences of the lookahead term on the next token through our choice of steering potential. The main lesson from this analysis is that the best choice, whenever possible, is one where the dependence of the potential $S$ on the token $a_i$ is concentrated on the term $s_i(\mathbf{a}_{\le i})$ of the decomposition, while future contributions $s_j(\mathbf{a}_{\le j})$ for $j > i$ have as small a dependence as possible on previous tokens $a_i$. This has been the guiding principle behind the steering potentials we have employed in the computational experiments of this work.

### G. Computational complexity of ILMC

Here we derive the asymptotic computational complexity of the ILMC algorithm with respect to the total sequence length $L$. We assume that the primary computational bottleneck is the evaluation of the generative energy $E_t(\mathbf{a}_{\le t})$, which requires a forward pass of the Transformer model. The cost of computing the steering potential $S_t(\mathbf{a}_{\le t})$ is assumed to be negligible in comparison.

In a standard Transformer-based autoregressive model with key-value caching, generating the $t$-th token requires computing attention over the previous $t-1$ tokens. The computational cost for a single forward pass at step $t$ is therefore proportional to $t$. The total cost to generate a sequence of length $L$ is:

$$\mathcal{C}_{\text{gen}} \propto \sum_{t=1}^{L} t \approx \frac{1}{2}L^2 \implies O(L^2) \tag{69}$$

In our proposed method, the sequence grows in blocks of size $B$. Let $M = L/B$ be the total number of elongation steps. We index these steps by $n = 1, \ldots, M$. At step $n$, the current sequence length is $t_n = n \cdot B$.

The computational cost at step $n$, denoted $C(n)$, consists of two components: the elongation phase, where $B$ new tokens are generated, and the refinement phase, which involves performing $N_{\text{MC}}$ resampling steps over a window of size $K$.

For the refinement step at length $t_n$, we perform $N_{\text{MC}}$ iterations. In each iteration, we resample a suffix of length roughly proportional to $K$ (on average $K/2$). Since the current sequence length is $t_n$, the cost to regenerate these tokens is proportional to the context length multiplied by the number of tokens generated: $K \cdot t_n$. Thus, the cost of one full refinement phase at step $n$ is:

$$\mathcal{C}_{\text{ref}}(n) \propto N_{\text{MC}} \cdot K \cdot t_n \tag{70}$$

The total cost $\mathcal{C}_{\text{total}}$ is the sum over all elongation steps $n$:

$$\begin{aligned}\mathcal{C}_{\text{total}} &\approx \sum_{n=1}^{L/B} (\mathcal{C}_{\text{elong}}(n) + \mathcal{C}_{\text{ref}}(n)) \\ &\approx \sum_{n=1}^{L/B} N_{\text{MC}} K(nB) \quad \text{(Dominant term)}\end{aligned} \tag{71}$$

Factoring out constants and substituting the sum of integers $\sum_{n=1}^{M} n \approx \frac{1}{2}M^2$:

$$\mathcal{C}_{\text{total}} \propto N_{\text{MC}} KB \sum_{n=1}^{L/B} n \approx N_{\text{MC}} KB \frac{1}{2} \left(\frac{L}{B}\right)^2 \tag{72}$$

Simplifying this expression yields:

$$\mathcal{C}_{total} \propto \frac{N_{\text{MC}} K}{2B} L^2 \tag{73}$$

From the derived expression, we identify two distinct scaling regimes based on the choice of the lookback window $K$.

- First, in the case of non-extensive lookback ($K = \text{const}$), if $K$ is a fixed hyperparameter independent of the sequence length, then $K$ acts as a constant prefactor. The scaling becomes:

$$\mathcal{C}_{\text{total}} \propto O(L^2) \tag{74}$$

  This matches the complexity class of standard autoregressive sampling.

- Second, in the case of extensive lookback ($K \propto L$), if the lookback window grows with the sequence length (e.g., $K = L$ to ensure global optimality as per Proposition 1), we substitute $K \propto L$ into the equation:

$$\mathcal{C}_{\text{total}} \propto L \cdot L^2 \implies O(L^3) \tag{75}$$

  In this regime, the algorithm incurs a cubic computational cost.

### H. Model Architecture and Training

To validate ILMC, we used two autoregressive Transformer architectures on the Chorismate Mutase (CM) and Phage Lysozyme (PL) families. In addition, for the classifier-guided steering experiments on the Response Regulator (RR) family, we trained an additional sDO.

The first architecture is ProGen3 (Bhatnagar et al., 2025), a family of eight autoregressive pLMs that employ a sparse mixture of experts (MoE) transformer decoder architecture. These models range in scale from 112M to 46B parameters and were pre-trained on a massive corpus of 3.4 billion protein sequences. For our experiments, we fine-tuned the smallest ProGen3 version (ProGen3-112M) on the Chorismate Mutase and Phage Lysozyme families, as we found it to be already complex enough to correctly model the related sequence landscapes. This version of the model features 10 MoE layers with a latent dimension of 384, 6 attention heads, 8 experts per layer, RMSNorm for layer normalization, SiLU activation function, and rotary positional embedding with a context length up to 65,536 tokens. For both families, we reserved a random subset of 1000 sequences as a validation set, keeping 17,094 training sequences for Chorismate Mutase and 10,859 training sequences for Phage Lysozyme, and selected the same set of parameters for fine-tuning, without further optimization. In particular, we set the learning rate to $5 \cdot 10^{-4}$, the batch size to 512, and trained the model until minimization of the validation loss, evaluated at the end of each epoch, with an early stopping patience of 3 epochs. We minimized the same pre-training loss function, obtained as the sum of the autoregressive loss (with weight 1.00) and the MoE load balancing loss (with weight 0.05). The procedure resulted in 5 fine-tuning epochs (170 steps) for Chorismate Mutase and 6 (132 steps) for Phage Lysozyme.

The second architecture is a custom, lightweight Decoder-only Transformer (sDO) trained from scratch using the exact same training and validation splits described above (17,094 training sequences for CM, 10,859 for PL and 855,395 aligned sequences for RR). This model comprises approximately 0.8 million parameters and features a context size sufficient to cover the full length of the sequences, which were short enough to avoid truncation. The architecture consists of 16 attention blocks with an embedding dimension of 64. Each block employs a Multi-Head Attention mechanism with 16 heads ($d_{head} = 4$). The sDO utilizes Rotary Positional Embeddings (RoPE), applies Layer Normalization before the attention and MLP blocks, and uses the GeLU activation function. For both families, the sDO was trained with a batch size of 512 and a learning rate of $10^{-3}$ using the AdamW optimizer. We employed a validation loss minimization strategy, which resulted in convergence at iteration 7,500 for Chorismate Mutase, iteration 4,500 for Phage Lysozyme and iteration 12'500 for Response Regulator.

For both architectures, the training sequences were curated by using Hidden Markov Models (HMM) to scan the UniProt database (The UniProt Consortium, 2025). For the Chorismate Mutase family, we constructed an HMM based on natural data from (Russ et al., 2020), while for Phage Lysozyme and Response Regulator, we utilized the standard Pfam HMM profile (Mistry et al., 2021). All entries containing more than 15% gaps in the resulting Infernal (Nawrocki & Eddy, 2013) alignment were removed, and the resulting datasets were used in their unaligned form.

### I. Melting Temperature Prediction

Predicted melting temperatures ($T_m$) were estimated using TemStaPro (Pudžiuvelytė et al., 2024), a tool based on ProtT5-XL (Elnaggar et al., 2022) embeddings that classifies sequence stability across thresholds from $40°\text{C}$ to $65°\text{C}$. To compute the statistics reported in Table 1, we mapped the predicted stability bins to scalar values using the bin centers. Specifically, sequences predicted as unstable at $40°\text{C}$ were assigned a value of $37.5°\text{C}$, those stable at $65°\text{C}$ were assigned $67.5°\text{C}$, and all intermediate bins were assigned their arithmetic midpoint (e.g., stability in the $[40°\text{C}, 45°\text{C})$ range was assigned $42.5°\text{C}$).

### J. Steering potential for thermostability enhancement

The amino acid composition of proteins in thermophilic organisms differs systematically from that of mesophilic organisms. Specifically, thermophilic proteomes show a marked enrichment in charged residues (such as Glu, Arg, and Lys) and a corresponding depletion of polar uncharged residues (such as Gln and Asn). These compositional adaptations facilitate the formation of stabilizing interactions, such as salt bridges, and reduce the presence of residues prone to thermal deamidation (Zeldovich et al., 2007; Fukuchi & Nishikawa, 2001).

To construct our steering potential, we utilized the ratio of amino acid frequencies observed on the surfaces of thermophilic proteins relative to mesophilic proteins (Fukuchi & Nishikawa, 2001). We relied specifically on surface composition rather than whole-chain metrics because the protein interior is strictly constrained by structural packing requirements, whereas surface residues possess greater evolutionary freedom to adapt to environmental stressors. To center the potential such that a score of zero represents a neutral mesophilic background, we define the weight $w(a)$ for amino acid $k$ as:

$$w(a) = \frac{f_{\text{thermo}}(a)}{f_{\text{meso}}(a)} - 1 \tag{76}$$

Positive values indicate residues associated with thermostability, while negative values indicate residues associated with mesophilic environments. The steering function $S(\mathbf{a})$ for a sequence $\mathbf{a} = (a_1, \dots, a_L)$ is defined as the negative sum of these weights:

$$S^{\text{Tm}}(\mathbf{a}) = -\sum_{i=1}^{L} w(a_i) \tag{77}$$

The complete set of weights used in our experiments is provided in Table 6.

Figure 12 illustrates the scatter plot of predicted melting temperatures against the defined thermophilic steering score for the natural sequences. While the correlation is robust across the generated designs, it appears less pronounced in the natural sequences. This is primarily because the natural sequences for Chorismate Mutase and Phage Lysozyme are predominantly mesophilic, clustering within the low melting temperature regime ($T_m < 40°\text{C}$). Also, the natural data exhibits limited variability in the thermophilic amino acid scores relative to the broader landscape explored by our sampling method.

*Table 6.* Thermophilic propensity weights derived from surface amino acid ratios. Weights are calculated as (Ratio $-$ 1), centered at 0.

| CATEGORY | AMINO ACID | RATIO | WEIGHT $w(a)$ |
|---|---|---|---|
| **STRONGLY ENRICHED** | GLU (E) | 1.28 | +0.28 |
| (RATIO $>$ 1.10) | LYS (K) | 1.27 | +0.27 |
| | TRP (W) | 1.24 | +0.24 |
| | ARG (R) | 1.23 | +0.23 |
| | TYR (Y) | 1.22 | +0.22 |
| | ILE (I) | 1.21 | +0.21 |
| | PHE (F) | 1.21 | +0.21 |
| | LEU (L) | 1.14 | +0.14 |
| | VAL (V) | 1.13 | +0.13 |
| | PRO (P) | 1.13 | +0.13 |
| **NEUTRAL** | GLY (G) | 1.04 | +0.04 |
| (0.90 $-$ 1.10) | CYS (C) | 0.94 | -0.06 |
| | MET (M) | 0.91 | -0.09 |
| **STRONGLY DEPLETED** | ASP (D) | 0.82 | -0.18 |
| (RATIO $<$ 0.90) | SER (S) | 0.79 | -0.21 |
| | ASN (N) | 0.71 | -0.29 |
| | THR (T) | 0.71 | -0.29 |
| | ALA (A) | 0.66 | -0.34 |
| | HIS (H) | 0.62 | -0.38 |
| | GLN (Q) | 0.55 | -0.45 |

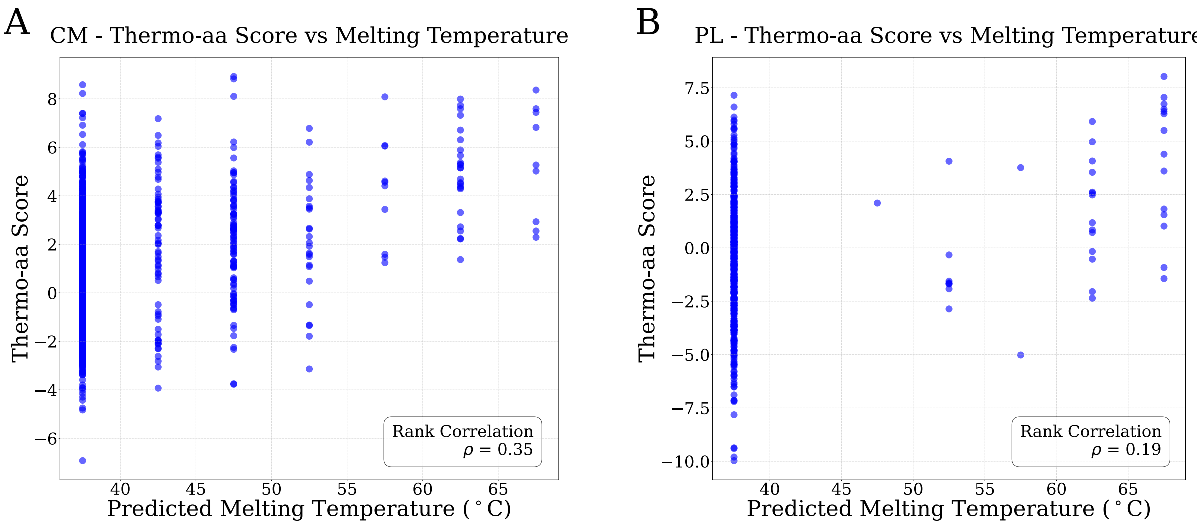


*Figure 12.* **Correlation between thermophilic amino acid score $-S^{\mathrm{Tm}}(a)$ and predicted melting temperature.** Scatter plots showing the relationship between the steering score (Thermo-aa Score) and predicted melting temperature ($T_m$) for 1024 natural sequences. **(A)** Chorismate Mutase (CM) family (Rank Correlation $\rho = 0.35$). **(B)** Phage Lysozyme (PL) family (Rank Correlation $\rho = 0.19$). The observed correlations are limited primarily because the natural sequences are predominantly mesophilic, clustering within the lowest temperature bin ($T_m < 40°$C).

### K. Steering potential for sampling in the vicinity of a wild-type sequence

The necessity of the modified definition for the steering potential $S_t^{\mathrm{WT}}(\mathbf{a}_{\leq t})$ arises from the inability of the standard edit distance to provide a meaningful steering signal during the early stages of autoregressive generation.

Consider the standard edit distance $\mathrm{EditDistance}(\mathbf{x}, \mathbf{y})$. If we were to naively use this as our partial steering potential $S_t^{\mathrm{WT}}(\mathbf{a}_{\leq t}) = \mathrm{EditDistance}(\mathbf{a}_{\leq t}, \mathbf{w})$ against a target wild-type $\mathbf{w}$, the steering signal $S_t^{\mathrm{WT}}(\mathbf{a}_{\leq t})$ would remain practically constant for any prefix length $t \ll L_w$. This occurs because, in most cases, the edit distance is entirely determined by the length difference between the target $\mathbf{w}$ and the partial generation $\mathbf{a}_{\leq t}$, rendering the specific identity of the generated tokens irrelevant.

This failure is best illustrated through an example. Let the target protein be:

$$\mathbf{w} = \mathrm{KKKKKKDNRCQA} \tag{78}$$

Assume the model has correctly generated the prefix $\mathbf{a}_{<t} = \mathrm{KKK}$. We now consider the emission of the next token $a_t$:

- Scenario A (Correct Continuation): We emit $a_t = \mathrm{K}$, forming the prefix KKKK.
- Scenario B (Incorrect Continuation): We emit $a_t = \mathrm{A}$, forming the prefix KKKA.

Intuitively, Scenario A should be highly preferred as it matches the local sequence context. However, under the standard edit distance against the full target $\mathbf{w}$, both scenarios yield an identical distance score:

$$\begin{aligned} &\mathrm{EditDistance}(\mathrm{KKKK}, \mathbf{w}) = 9 \quad \text{(9 deletions to match)} \\ &\mathrm{EditDistance}(\mathrm{KKKA}, \mathbf{w}) = 9 \quad \text{(Match final 'A', delete intermediate)} \end{aligned} \tag{79}$$

In Scenario B, the algorithm matches the newly emitted 'A' to the final 'A' of the target string $\mathbf{w}$, reaching the exact same distance because the cost of deleting the ending portion of the target is identical to the cost of deleting internal segments. Practically, for a prefix $\mathbf{a}_{\leq t}$ where $t \ll L_w$, the distance is determined by the number of deletions required to account for the length difference, since almost any amino acid generated at step $t$ is likely to appear somewhere later in a long target chain. Consequently, the sampler receives no signal indicating whether the generated token helps in steering the generation toward the target sequence.

Our proposed definition, $S_t^{\mathrm{WT}}(\mathbf{a}_{\leq t}) = \min_k \mathrm{EditDistance}(\mathbf{a}_{\leq t}, \mathbf{w}_{\leq k})$, resolves this problem by effectively eliminating the penalty for deleting the ending part of the target:

- For KKKK, we compare against all prefixes of $\mathbf{w}$. The best match is $\mathbf{w}_{\leq 4}$ ("KKKK"), yielding a distance of $0$.
- For KKKA, we compare against all prefixes. The best match (not unique) is against $\mathbf{w}_{\leq 4}$ ("KKKK"), where 'A' results in a mismatch, yielding a distance of $1$.

This metric successfully creates a steering signal that can accurately inform the sampling regarding the similarity with the target sequence.

### L. Acceptance Rate Decay

The original sampling framework proposed by (Karan & Du, 2025) for text-based architectures is designed to sample from the exponentiated distribution of the model itself, $P(\mathbf{a})^\beta$. In that unsteered scenario, the proposal distribution—standard autoregressive elongation at inverse temperature $\beta$—is naturally aligned with the target distribution. Consequently, the acceptance rates remain relatively high even for larger lookback windows, as the proposal provides a good approximation of the target.

The introduction of an external steering potential $S(\mathbf{a})$ creates a discrepancy between the proposal distribution and the target distribution. In ILMC framework, the proposal steps (Eq. 4) are generated via the base model, which is agnostic to the steering potential $S(\mathbf{a})$. The steering constraint is enforced solely through the acceptance step of the Metropolis-Hastings transition:

$$A(\mathbf{a} \to \mathbf{a}') = \min\left(1, e^{-\beta\Delta E - \lambda\Delta S}\,\frac{q(\mathbf{a})}{q(\mathbf{a}')}\right) \tag{80}$$

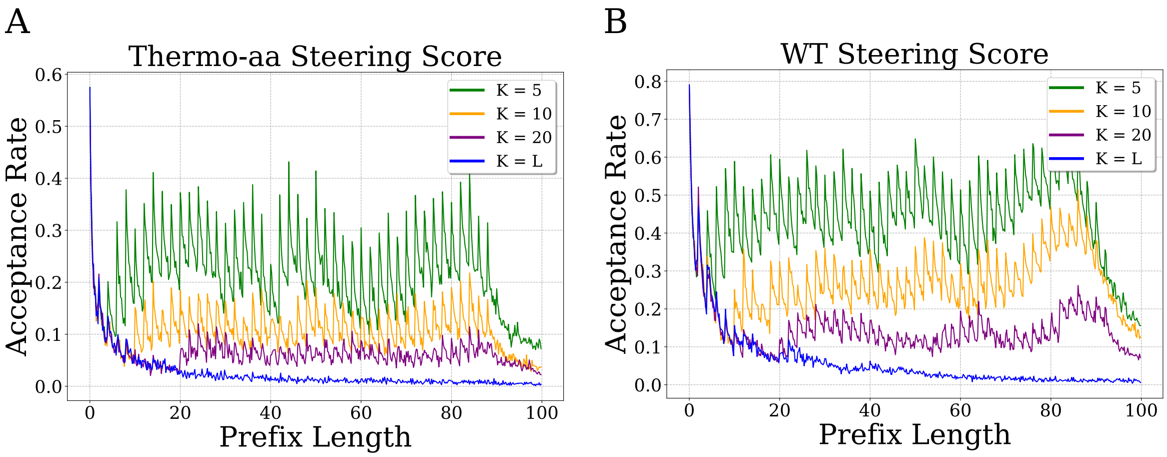


*Figure 13.* **Metropolis acceptance rate as a function of the generated prefix length** during ILMC for the sDO architecture on the CM family ($B = 2$, $N_{\text{MC}} = 10$, averaged over 1024 sequences). For both steering with $S^{Tm}$ (A) and $S^{WT}$ (B), we observe a sharp decline in acceptance rate as the lookback window $K$ increases. This indicates that while larger $K$ brings the distribution closer to sampling from $Q^*$, it renders the sampling procedure significantly less efficient.

where $\Delta E = E_t(\mathbf{a}'_{\leq t}) - E_t(\mathbf{a}_{\leq t})$ and $\Delta S = S_t(\mathbf{a}'_{\leq t}) - S_t(\mathbf{a}_{\leq t})$. When the lookback window $K$ is large, the proposal mechanism effectively generates a long sequence segment "blindly," without conditioning on the steering objective. The probability that such an unguided trajectory spontaneously satisfies the target steering constraints decays with the length of the segment. Consequently, newly proposed states frequently result in unfavorable steering shifts (large $\Delta S$), causing the term $e^{-\lambda \Delta S}$ to go to 0, leading to high rejection rates.

To mitigate this issue and ensure computational efficiency, it is practically necessary to restrict the size of the lookback window $K$. By using a shorter window, the proposal trajectory is forced to remain in the vicinity of the existing prefix, which has already been steered into a favorable region of the landscape, thereby increasing the probability of acceptance. We observe empirically that the acceptance rate is inversely related to $K$ (Figure 13). To conclude, while $K = L$ is theoretically required to sample exactly from $Q^*$, a restricted $K$ is practically required to maintain a feasible acceptance rate when the steering strength $\lambda > 0$.

### M. Pareto Frontier of Generative Quality and Steering Potential

When sampling from the target distribution $Q^*(\mathbf{a}) \propto \exp(-\beta E(\mathbf{a}) - \lambda^{\text{Tm}} S^{\text{Tm}}(\mathbf{a}))$, the objective is to identify sequences that simultaneously minimize the generative energy $E(\mathbf{a})$, ensuring biological viability, and the steering potential $S^{\text{Tm}}(\mathbf{a})$, reflecting the desired trait. However, because the steering potential is not perfectly aligned with the natural sequence distribution learned by the base model, these two objectives are likely to compete. Optimizing for the steering potential beyond a certain threshold requires the sampler to explore regions of the sequence space that the model considers less probable, thereby increasing the generative energy. This competition defines a Pareto frontier in the $(E, S)$ plane, representing the set of optimal trade-offs where one metric cannot be improved without degrading the other. To empirically illustrate this relationship, we performed a systematic analysis with the sDO base model, using the thermophilic amino acid score Eq. (8) as the steering potential by varying the inverse temperature $\beta$ and the steering coefficient $\lambda^{\text{Tm}}$. As shown in Figure 14, the resulting samples clearly delineate a frontier; it is not possible to achieve high steering scores without sacrificing generative quality, and vice versa.

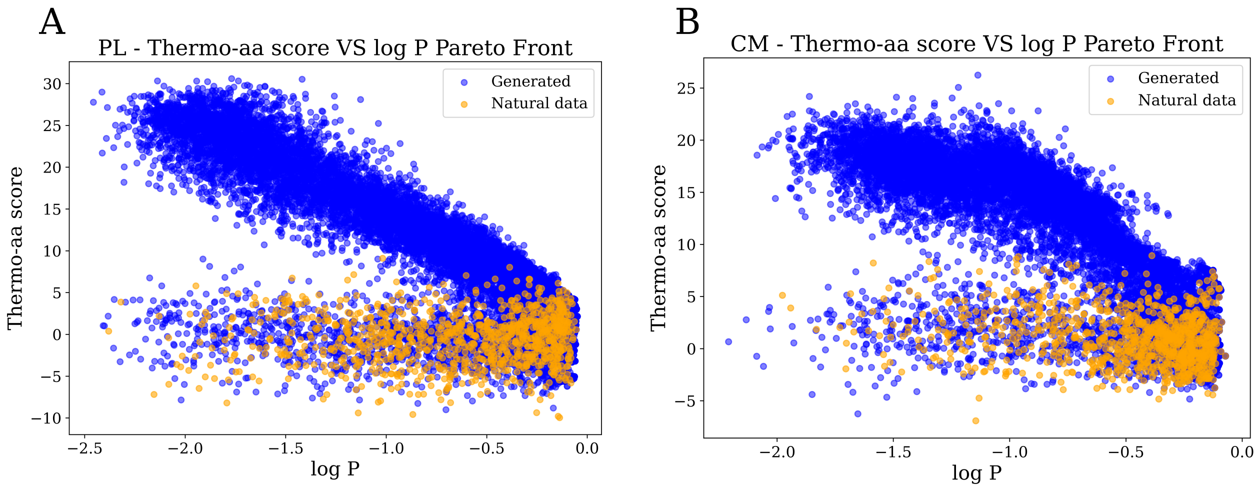


*Figure 14.* The Pareto frontier between the mean per-token log-probability ($\log P$) and the thermophilic amino acid score $-S$ for the sDO models trained on Phage Lysozyme (PL) and Chorismate Mutase (CM). For each protein family, libraries of 1024 sequences were sampled utilizing ILMC with fixed parameters $B = 2$, $K = 5$, $N_{\mathrm{MC}} = 10$, across a dense grid of inverse temperature $\beta$ and steering strength $\lambda^{\mathrm{Tm}}$. For CM, the grid parameters were $\beta \in \{1, 1.8, 2.6\}$ and $\lambda^{\mathrm{Tm}} \in \{0, 5, 10, 15, 20\}$; for PL, $\beta \in \{1, 1.3, 2\}$ and $\lambda^{\mathrm{Tm}} \in \{0, 5, 10, 15, 20\}$. Natural sequences are indicated in orange for reference. The plots clearly demonstrate the Pareto frontier in the upper-right corner, illustrating the intrinsic trade-off between maximizing generative likelihood and enforcing the targeted steering potential.

### N. Double Steering

Another interesting application of ILMC is the simultaneous imposition of multiple steering potentials. For instance, we could aim to enhance thermostability while strictly maintaining the generation within the mutational vicinity of a known wild-type. Mathematically, the extension of our method to two objectives is straightforward. The target distribution becomes:

$$Q^*(\mathbf{a}) \propto \exp\left(-\beta E(\mathbf{a}) - \lambda^{\mathrm{WT}} S^{\mathrm{WT}}(\mathbf{a}_{\leq L}) - \lambda^{\mathrm{Tm}} S^{\mathrm{Tm}}(\mathbf{a})\right) \tag{81}$$

To demonstrate the efficacy of this approach, we targeted a specific CM wild-type sequence (Index 2 in the dataset), chosen for its low predicted melting temperature ($T_m < 40$). Our objective is to generate variants possessing between 14 and 16 mutations relative to this wild-type, while simultaneously pushing for higher thermostability.

We compared two generation strategies:

1. Distance-Only Steering: We sampled using parameters tuned to target the 14-16 mutation range without any thermo-aa composition bias ($\beta = 1.0, \lambda^{\mathrm{WT}} = 7.5, \lambda^{\mathrm{Tm}} = 0$).
2. Double Steering: We sampled using parameters tuned to target the same mutational range but with thermo-aa composition bias ($\beta = 1.3, \lambda^{\mathrm{WT}} = 6.5, \lambda^{\mathrm{Tm}} = 6.5$).

From the resulting $2048$ generated sequences, we selected the ones that met two criteria: (1) exactly 14 to 16 edit distance from the wild-type, and (2) a generative log-likelihood greater than the average likelihood of unsteered autoregressive samples $\log P(\mathbf{a}) > -0.8$. This ensures that we compare only high-quality, viable protein candidates.

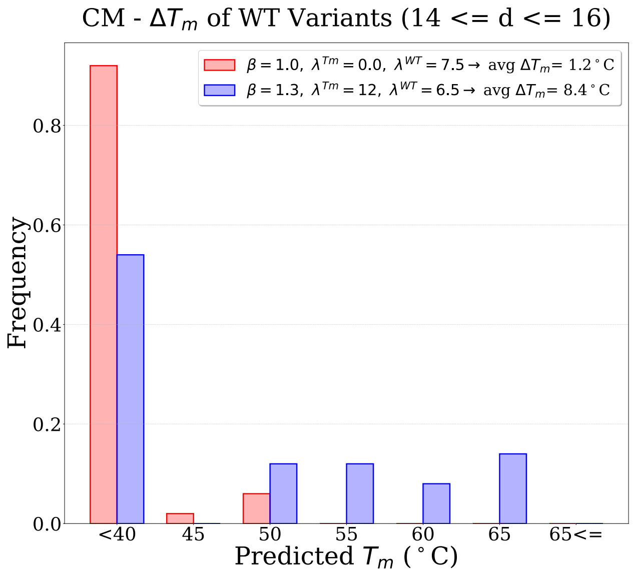


*Figure 15.* Distribution of predicted melting temperatures ($T_m$) for Chorismate Mutase (CM) variants generated within a 14–16 mutation shell relative to the target wild-type (WT $T_m = 37.5°$C). Sequences were sampled using the sDO architecture via ILMC ($B = 2, K = 10, N_{\mathrm{MC}} = 100$). We compare two strategies: "Distance Only" ($\beta = 1.0, \lambda^{\mathrm{WT}} = 7.5, \lambda^{\mathrm{Tm}} = 0$) and "Double Steering" ($\beta = 1.3, \lambda^{\mathrm{WT}} = 6.5, \lambda^{\mathrm{Tm}} = 6.5$), with all candidates filtered for high likelihood ($\log P > -0.8$). The figure displays the top 50 sequences ranked by thermophilic amino acid score $-S^{\mathrm{Tm}}(\mathbf{a})$ from total populations of 185 (Distance Only) and 155 (Double Steering) variants, demonstrating that double steering successfully shifts the distribution toward higher $T_m$ while maintaining the target edit distance.

As illustrated in Figure 15, the multi-objective approach proves effective: the average $\Delta T_m$ increases from $1.2°$C in the distance-only baseline to $8.4°$C) in the double-steered case, demonstrating that ILMC can effectively navigate the intersection of multiple constraints to design stable, high-likelihood mutations.

The practical challenge lies in the hyperparameter search. Balancing the inverse temperature $\beta$, the distance penalty $\lambda^{\mathrm{WT}}$, and the thermo-bias $\lambda^{\mathrm{Tm}}$ requires a non-trivial search to ensure the sampler explores the correct intersection of the energy landscape (high likelihood, specific distance, high stability).

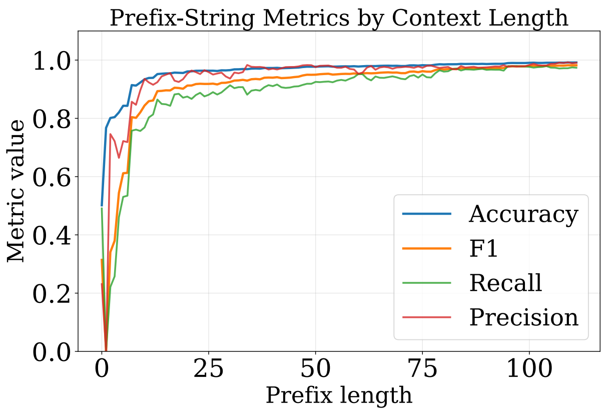


*Figure 16.* **Any-prefix classifier metrics versus prefix length** for the Response Regulator benchmark. Test-time accuracy, precision, recall, and F1 are reported as a function of the available prefix length.

### O. Any-prefix classifier for Response Regulators

For classifier-guided steering on the Response Regulator (RR) family, we trained a lightweight any-prefix classifier to predict the target GerE homodimerization class from incomplete sequence prefixes. The classifier was trained on the same aligned RR dataset used for the generation experiments, comprising 855,395 sequences of length 111, of which 23.26% belong to the positive class.

The model receives as input a prefix $\mathbf{a}_{\leq t}$, padded to length 111, and outputs a binary logit. Architecturally, it consists of a token embedding layer with vocabulary size 21 and embedding dimension 16, followed by flattening of the full embedded prefix, a fully connected layer from 1776 to 16 units, a ReLU activation, and a final linear layer from 16 to 1. The total number of trainable parameters is 28,785.

We trained the classifier with weighted binary cross-entropy to account for class imbalance, sampling prefix lengths uniformly during training so that the model learned to score arbitrary contexts rather than only full sequences. Training required approximately 5 minutes on a single NVIDIA RTX 4090 GPU.

Classifier performance was then evaluated as a function of prefix length. As expected, very short prefixes are only weakly informative, whereas predictive performance improves steadily as longer prefixes are observed. The complete dependence of the evaluation metrics on prefix length is reported in Fig. 16.

### P. Hyperparameter Selection Guide

A few practical heuristics can be used to initialize the ILMC hyperparameters $(\beta, \lambda, K, N_{\mathrm{MC}})$.

Selection of $\beta$: In practice, we first tune $\beta$ in the unsteered setting ($\lambda = 0$) so that the generated sequences match a desired mean per-token $\log P$. A useful reference point can be obtained from the distribution of $\log P$ on natural sequences; in particular, the empirical mean over the natural data provides a convenient starting target.

Selection of $\lambda$: A useful rule of thumb for choosing $\lambda$ is to set it such that, for typical local proposals, the steering contribution is of the same order as the generative contribution $\beta\Delta E \sim \lambda\Delta S$. This prevents either term from dominating the Metropolis acceptance ratio.

Selection of $K$: In our experiments, moderate lookback windows were usually sufficient, and values of $K$ between 5 and 10 already yielded strong improvements over purely autoregressive sampling. Larger values of $K$ can be useful when maximizing diversity is especially important, but in practice they often reduce the acceptance probability substantially and therefore typically require larger $N_{\mathrm{MC}}$, especially in the steered setting.

Selection of $N_{\mathrm{MC}}$: The number of refinement steps $N_{\mathrm{MC}}$ should then be increased until the chain performs a non-trivial number of accepted corrections during generation. Empirically, we found it useful to tune $N_{\mathrm{MC}}$ until acceptance rates during elongation were on the order of $\sim 0.3$. It is also useful to monitor the acceptance rate as a function of the refinement step: if it still fluctuates strongly across the $N_{\mathrm{MC}}$ updates, this suggests that a larger value may be needed to approach a local equilibrium over the current prefix. Different steering potentials require different values of $N_{\mathrm{MC}}$ depending on their ruggedness. For the thermostability steering potential, relatively small values of $N_{\mathrm{MC}}$ were sufficient, because the score is additive and each token swap produces a small graded variation in $S_t^{\mathrm{Tm}}$. By contrast, the wild-type proximity objective required substantially larger $N_{\mathrm{MC}}$, since most local proposals only weakly affect the distance-based score and many accepted moves are needed to accumulate a significant shift toward the target sequence.

Overall, we found the following initialization strategy to work well in practice: tune $\beta$ to match the target mean $\log P$, set $\lambda$ so that $\lambda \Delta S \sim \beta \Delta E$ for typical mutations, use $K \in [5, 10]$, and increase $N_{\mathrm{MC}}$ until acceptance rates during generation are around $0.3$.

Because ILMC operates sequentially, hyperparameter search can be accelerated by working on partial sequence prefixes of about 20–30 tokens. In practice, one can generate only the first 20–30 residues and monitor acceptance rate, mean $\log P$, and steering score; this already provides a useful proxy for the behavior of the full sampler while significantly reducing computational cost.
